## Supplementary Material for "Knowledge synthesis from 100 million biomedical documents augments the deep expression profiling of coronavirus receptors"

1. nference, inc., One Main Street, Suite 400, East Arcade, Cambridge, MA 02142, USA.
2. nference Labs, pvt. ltd., Wind Tunnel Road, Murugesh Palaya, Bengaluru 560017, India.
3. Janssen Research & Development LLC, USA.
4. nference, 111 Peter St, Toronto, ON M5V 2G9, Canada.

#### Evaluation of literature-derived association scores

**Ground truth** We need some notion of ground truth to evaluate the quality of association measures. We use sets of known pairs of related entities versus a "control" group of random pairs of entities of the same classes. We use a few different sets of known pairs:

(1) Disease-Gene relationships based on OMIM[1]
(2) Drug-Gene relationships
(3) Drug-Disease relationships based on fda labels
  (a) Drugs and their on-label indications
  (b) Drugs and their on-label adverse events
(4) Logical queries for ambiguous tokens

One demonstration of the use of the logical query system is to disambiguate a token by conjoining it with a disambiguating token. An example is clearer: the token "egfr" can refer to the gene entity *epidermal growth factor receptor*, but also the test measure entity *estimated glomerular filtration rate*. A query "egfr AND kidney" should return results related to the latter meaning, while "egfr AND lung_cancer" the former. In particular, an unambiguous referent to the right entity should be highly related to the query. So example known pairs in this data are ("egfr AND kidney", "estimated_glomerular_filtration_rate") and ("egfr AND lung_cancer", "epidermal_growth_factor_receptor"). We used an internal set of a ~200-300 such ("A AND B", "C") pairs (originally built up for other reasons).

Note: One key drawback of the word2vec vector cosine similarity[1,2] method is its inability to get scores for logical queries as described above, because the method[2] does not address the question of how to get vectors for queries that are logical combinations of tokens.

**Evaluation metrics**
Given a scoring method and a particular set of positive/control pairs, we get two sets of scores: one set for the positive pairs and one set for the negative pairs.

*Cohen's d* - We compute the *Cohen's d* standard statistical measure of distance between 2 samples [3].

*Mann-Whitney U (normalized)* The Mann-Whitney U is a nonparametric measure of distribution distance: it counts the number of transposed pairs[4].

*Metrics based on training a 1-d logistic model*

In this test we are discriminating between 2 classes (true association/non-association) based on 1 feature. We have two metrics based on fitting a 1-feature logistic curve to the data. (**Figures S1A-B)**

*Brier score:* The Brier score is the average squared error of the logistic curve above: that is, for each labeled point, we square the vertical distance to the logistic curve, and average over all labeled points[5].

*Log loss*[6] The logistic log loss is the average *-log [model probability of true label]* for each labeled point. If the model is perfect at the point, it incurs no loss. If it predicts 0.5, it incurs -*log[0.5]* loss. If it predicts "yes" with certainty when the answer is "no" it incurs infinite loss (a logistic function never touches 0 or 1 so this won't happen in our case).

*Neg log percentile*: For most of the scoring rules, we also include a *-log(percentile)* version of the rule. This is constructed as follows, for query *q*, token *t*, and score *S(q, t)*

1) Compute the scores *S(q, t')* for *q* with every token *t'*. Let *R* be the number of these that are nonzero.
2) Take the rank *r* of *S(q, t)* among all nonzero *S(q, t')*.
3) The *neg log percentile* score *nlS(q, t)* associated with *S* is *-log(r/R)*

We do this to
(i) control for differences across queries
(ii) control for differences in the shapes of the distributions that different association scoring functions take.

This procedure maps all the *S(q, t')* to an Exponential(1) distribution. We chose Exponential(1) because it is simple, intuitively reasonable and many of the scores naturally seemed to be approximately exponential.

**Results of evaluation**

The table below lists the performance approximately 2100 disease-gene pairs.

| Assoc score↓ | Cohen's d (+) | Mann-W U norm. (-) | Logistic log loss (-) | Logistic Brier score (-) |
|---|---|---|---|---|
| Cosine (w2v) | 1.31 | 0.197 | 0.51 | 0.168 |
| Raw PMI | 2.07 | 0.0953 | 0.374 | 0.116 |
| Raw PMI -log(pctile) | 2.15 | 0.0947 | 0.355 | 0.111 |
| Exp PMI | 2.17 | 0.0897 | 0.356 | 0.109 |
| Exp PMI -log(pctile) | 2.21 | 0.0903 | 0.341 | 0.105 |
| Raw Local Score | 2.35 | 0.0828 | 0.312 | 0.0947 |
| Raw Local Score -log(pctile) | 2.28 | 0.0832 | 0.317 | 0.0963 |
| Exp Local Score | 2.34 | 0.0812 | ***0.301** | ***0.0915** |

| | | | | |
|---|---|---|---|---|
| Exp Local Score -log(pctile) | ***2.36** | ***0.0811** | 0.308 | 0.093 |
| log(coocc) | 2.24 | 0.097 | 0.348 | 0.105 |

*Interpretation of the above table*

Each row corresponds to an association score whereas each column corresponds to one of the evaluation metrics. A (+) in the column means a higher evaluation metric value, the better the association score in that row separates the positive and random pairs. A (-) mean a lower evaluation metric is better. Note all the metrics are immune to linear rescalings; also the Mann-Whitney U score is nonparametric.

**High-dimensional Word Embeddings for determining the significant global associations**

**Figure S1C** illustrates two histograms generated from a random set of vectors (in the vector space generated by the Neural Network) where one distribution (denoted as "DISTANCE < 0.32") represents all vector pairs whose cosine distance is less than 0.32 (deemed "not strong associations") and the other distribution (denoted as "DISTANCE >= 0.32") represents all vector pairs whose cosine distance is greater than 0.32 (deemed "strong associations"). This can show how common a phenomenon it is to find word vector pairs that have very good cosine distances but yet not co-occur even once in the corpus. The "DISTANCE >=0.32" bar at zero value suggests that roughly 11% of vector pairs whose cosine distances where greater than 0.32 ("*strong associations*") never occurred together even once in a document. It is also clear from the figure that albeit more of the mass of the "DISTANCE >=0.32" distribution is skewed to the right as expected (more co-occurrences and hence unsurprisingly larger cosine distances), there is a long tail of the "DISTANCE < 0.32" distribution (very high co-occurrences but small cosine distances). The long tail is a direct consequence of negative sampling—where vectors corresponding to common words that co-occur quite often with significant words in a sliding window are moved away from vectors of the other words.

**What does the word2vec neural network do? From the perspective of Genes-Diseases associations**

One way to view the word2vec "black box" operation from a Genes/Diseases perspective (cosine of <Gene, Disease> for all Genes and Diseases) as a Transfer Function which changed the input probability distribution (pre-training randomly assigned word vectors for Genes and Diseases) to a new probability distribution. The "null hypothesis" (which seems to be pretty much preserved in actuality in the way word2vec assigns random values to vectors) is the "green colored" Cosine Distribution (**Figure S1D**).

Once word2vec training is over, the final word vectors are placed in specific positions in the 300-dimensional space so as to present the "blue colored" Empirical distribution (the actual cosine distance between <Gene, Disease> pairs that we observe). The "orange curve" is the 2-Gamma mixture (the parametric distribution that in some sense captures the “empirical distribution” with just 8 parameters (2 alphas, 2 betas, 2 ts and 2 phis).

Few observations from this analysis:

- Note the “symmetrical” cosine distribution after training becomes “Asymmetrical” with a longer “right tail”. The asymmetry is the reason why Gamma distribution worked better than say, Gaussian, for the curve fit. The mean of the distribution gets shifted to the right after training as one would expect — the vectors during training are “brought together” by parallelogram addition most of the time explaining the shift to the right (negative sampling will cause a movement in the opposite direction but that will disproportionately affect the “ultra-high frequency” words — which to begin with get “more” positively sampled and hence the 3-gamma with a bump near 0.6 happens for ultra-high frequency words).
- The most interesting associations, by definition, are in the tail of the distribution.

**What does varying the number of dimensions in the word2vec space do to the underlying cosine distance distributions in a large textual corpora?**

**Figure S1E** illustrates a cosine distance probability density function (PDF) graph to visually describesthe implementation of the word2vec like Vector Space Model in various N-dimensional spaces. As described in the **Methods** section, the system is a Semantic Bio-Knowledge Graph of nodes representing the words/phrases chosen to be represented as vectors and edge weights determined by measures of Semantic Association Strength (e.g., the Cosine Distance between a pair of word embeddings represented as vectors in a large dimensional space). The cosine distance ranges from 0 (representing no semantic association) to 1 (representing strongest association). This metric of association can reflect the contextual similarity of the entities in the Biomedical Corpora. The typical dimensionality used by our neural network for generating the Global Scores is n=300 dimensions. This is because, as can be seen in the graph, the distribution is highly peaked with most of the mass centered around 0 -- that is, a randomly chosen pair of vectors typically are orthogonal or close to orthogonal, and furthermore over 300-dimensions, the distributions all have sufficiently long tails with the most interesting (salient) biomedical associations.

#### nferX Single Cell Platform Introduction

We have currently incorporated 32 human or mouse datasets (**Table S1**) which were manually selected to enable pan-tissue analyses such as the one presented here. We are in the process of scaling this platform to automatically consume all scRNA-seq datasets which are available in the public domain, and datasets from academic labs can be easily incorporated into the platform for public access by request. Currently, study data has been accessed in the form of counts matrices deposited into GEO or another public repository. These matrices have been uniformly processed using Seurat v3.1 including conversion into units of counts per 10,000 (CP10K), generation of

cluster assignments, and computation of UMAP and tSNE coordinates for dimensionality reduction plots. If cluster assignments were provided as associated metadata, we retain these assignments in the nferX platform; if cluster assignments were not provided, then clusters were manually annotated through an assessment of the cluster-defining genes[7].
To demonstrate the utility of the nferX Single Cell app, we briefly describe the use case of analyzing *Ins1* (insulin) expression in the Tabula Muris study[8] below. The platform contains three primary sections: (1) a summary of gene expression for a queried gene in individual cell populations (e.g., violin plots and summary statistics); (2) a two-dimensional visualization of all cell clusters from the selected study, along with overlaid gene expression data (dimensionality reduction plots); and (3) a summary of expression for all genes in every cell population identified in the selected study (e.g., cluster-defining genes).

When searching for *Ins1* expression in the Tabula Muris study, we first show the violin plot section which highlights that this gene is most highly expressed in pancreatic beta cells, with lower level detection in other cells derived from the pancreas. The cluster-level data is summarized within this table, including the number of cells with each cluster assignment, the percent of cells which express *Ins1* in each cluster, and the mean expression of *Ins1* in each cluster. The specificity of *Ins1* expression to any individual cluster is computed as a Cohen's D value, which measures the effect size between the mean expression in one cluster versus the mean expression in all other clusters. By default, clusters are shown in descending order of Cohen's D values; here as expected, pancreatic beta cells show the largest Cohen's D value for *Ins1* (**Figure S4A**).

On the left side of this table, the columns labeled as "Signals Strength to Tissue" and "Signals Strength to Cell" summarize the literature associations between *Ins1* and the tissue/cell combination in each row. Throughout the nferX platform, we use these "bubbles" to represent such literature associations, where the size of the bubble corresponds to the proximity of tokens throughout biomedical literature (e.g., how frequently does *Ins1* appear within 50 words of "pancreas"?), and the shading of the bubble corresponds to the similarity of tokens in a high-dimensional vector space of word embeddings (e.g., how similar is the "neighborhood" of words and phrases around *Ins1* compared to the neighborhood around "pancreas"?). Thus, bubbles which are large and dark indicate a concordant literature association in which these concepts are highly connected from both local and global perspectives.

In the next section, we display all single cells from the Tabula Muris in a dimensionality reduction plot, where UMAP is employed by default and tSNE can be selected as an alternative (**Figure S4B**). Note that this section is included for visualization purposes only, and these cluster visualizations were not used to assign cluster annotations. In the left panel, we display a "feature plot", in which the color of each point (i.e., cell) corresponds to the expression level of *Ins1* in that individual cell in units of ln(1+CP10K). In the right panel, cells are colored by a selected metadata feature; by default, cells are colored by tissue of origin for studies which contain multiple tissues and by annotated cell type for studies which only contain cells from one tissue.

In our final summary section, we summarize the genes which are most specifically expressed in each cell population from the selected study. By default, we prioritize those cell populations which were found to most strongly express the queried gene and display the cluster-defining genes in descending order by Cohen's D. Thus, here we show that the genes which are most specifically expressed in pancreatic beta cells include *Ins2, Iapp,* and *Ins1* (**Figure S4C**). In each row, we also compute the literature-derived associations between the gene and tissue/cell type to highlight well-known versus potentially underappreciated expression patterns.

##### List of studies included in the Single Cell Platform

The set of studies which have currently been analyzed and made accessible for analysis in the Single Cell Platform are listed below in **Table S1**.

| **Study Title** | **Tissue** | **Number of Cells** | **Number of Individuals** | **Species** |
|---|---|---|---|---|
| *Intra- and Inter-cellular Rewiring of the Human Colon during Ulcerative Colitis* | Colon | 110110 | 12 | Human |
| *A Cellular Anatomy of the Normal Adult Human Prostate and Prostatic Urethra.* | Prostate | 109061 | 3 | Human |
| *Single-cell reconstruction of the early maternal-fetal interface in humans* | Placenta | 64734 | 7 | Human |
| *Single-cell reconstruction of follicular remodeling in the human adult ovary* | Ovary | 57324 | 5 | Human |
| *Mature Kidney - Spatiotemporal immune zonation of the human kidney* | Kidney | 40268 | 8 | Human |
| *Census Of Immune Cells* | Bone Marrow | 38960 | 8 | Human |
| *Fetal Kidney - Spatiotemporal immune zonation of the human kidney* | Kidney | 27203 | 8 | Human |
| *Immune Cell Atlas: Blood Mononuclear Cells (2 donors, 2 sites)* | Blood | 24137 | 2 | Human |

| | | | | |
|---|---|---|---|---|
| *Single-cell transcriptomic atlas of the human retina identifies cell types associated with age-related macular degeneration* | Retina | 20091 | 3 | Human |
| *Single-cell reconstruction of the adult human heart during heart failure and recovery reveals the cellular landscape underlying cardiac function* | Heart | 14927 | 20 | Human |
| *Single-cell transcriptome analysis reveals differential nutrient absorption functions in human intestine* | Small intestine | 14537 | 2 | Human |
| *Spleen - Ischaemic sensitivity of human tissue by single cell RNA seq* | Spleen | 12049 | 4 | Human |
| *Esophagus - Ischaemic sensitivity of human tissue by single cell RNA seq* | Esophagus | 10810 | 3 | Human |
| *A human liver cell atlas reveals heterogeneity and epithelial progenitors.* | Liver | 10372 | 9 | Human |
| *A cellular census of human lungs identifies novel cell states in health and in asthma.* | Lung | 10235 | 4 | Human |
| *The adult human testis transcriptional cell atlas* | Testis | 6490 | 3 | Human |
| *Single-Cell Transcriptomic Analysis of Primary and Metastatic Tumor Ecosystems in Head and Neck Cancer* | HNSCC Tumor and Lymph Node | 5578 | 18 | Human |

| | | | | |
|---|---|---|---|---|
| *Identification of grade and origin specific cell populations in serous epithelial ovarian cancer by single cell RNA-seq* | Ovary | 3066 | 18 | Human |
| *Single-Cell Transcriptome Profiling of Human Pancreatic Islets in Health and Type 2 Diabetes.* | Pancreas | 2394 | 10 | Human |
| *A Single-Cell Transcriptome Atlas of the Human Pancreas.* | Pancreas | 2285 | 4 | Human |
| *De Novo Prediction of Stem Cell Identity using Single-Cell Transcriptome Data.* | Pancreas | 1004 | 5 | Human |
| *Transcriptome Landscape of Human Folliculogenesis Reveals Oocyte and Granulosa Cell Interactions.* | Ovary | 151 | 1 | Human |
| *Mapping the Mouse Cell Atlas by Microwell-Seq.* | 26 Tissues | 242533 | 69 | Mouse |
| *Single-cell transcriptomics of 20 mouse organs creates a Tabula Muris.* | 18 Tissues | 100605 | 132 | Mouse |
| *Single cell analysis reveals immune cell-adipocyte crosstalk regulating the transcription of thermogenic adipocytes* | Adipose | 36661 | 3 | Mouse |

| | | | | |
|---|---|---|---|---|
| *An atlas of the aging lung mapped by single cell transcriptomics and deep tissue proteomics* | Lung | 14813 | 15 | Mouse |
| *A single-cell survey of the small intestinal epithelium* | Small Intestine | 7216 | 6 | Mouse |
| *A revised airway epithelial hierarchy includes CFTR-expressing ionocytes* | Trachea | 7193 | 6 | Mouse |

##### *Expression of coronavirus receptors in other tissues by scRNA-seq*

We evaluated the expression profiles of ACE2, ANPEP, DPP4, and TMPRSS2 across various other human and mouse tissues by single cell RNA-sequencing. The complete set of tissues profiled is outlined in **Table S1** and includes both tissues which were identified from bulk RNA-sequencing as potential sites of expression for one or more receptors (e.g., kidney, adipose tissue, heart, testis, prostate, blood/immune system) as well as other tissues which did not show any evidence for expression at the bulk level (e.g., brain, retina, ovary, uterus). The results for several tissues are discussed in the Main text, including those of the respiratory and gastrointestinal tracts The analysis of data from all other tissues is discussed below with corresponding images corresponding found in **Figures S5-S20**.

Our analyses of bulk RNA-seq, proteomics, and IHC demonstrated that all coronavirus receptors are highly expressed in the kidney in addition to the small intestine (**Figures S3A-D, S5A-E**). scRNA-seq of human and murine kidney corroborates this finding, showing that ACE2, DPP4, and ANPEP are each robustly expressed in epithelial cells of the proximal tubules (**Figure S6**). Although clinical data suggests that SARS-CoV-2 does not reside in the urine (ref), we wondered whether ACE2, DPP4, and ANPEP show an overlapping pattern of expression heterogeneity among proximal tubular cells similar to that observed among small intestinal enterocytes. To address this, we computed cosine similarity scores between gene expression vectors specifically among ~27,000 recovered human proximal tubule epithelial cells[9] and found that again DPP4 and ANPEP are among the genes with the most similarly expression profile to ACE2 (**Figures S7A-C**). This is further supported by the strong transcriptional correlations among GTEx kidney samples (n = 85) by bulk RNA-seq, where DPP4 and ANPEP are again among the top 1% of gene correlations to ACE2 among the ~16,600 genes with mean expression > 1 TPM (**Figure S7D-F**). These observations are quite consistent with our previous analysis of small intestinal data and together may suggest an underlying transcriptional network which coordinates the expression of coronavirus entry receptors across diverse human tissues and cell types.

We then examined coronavirus receptor expression in the heart given the relatively high expression of ACE2 by bulk RNA-seq and the strong literature association identified between this gene-tissue pair (**Figure S8**). In the human heart, ACE2 is detected in 11% of cells or fewer from multiple populations including smooth muscle cells, cardiomyocytes, and fibroblasts. It is detected in a higher fraction of cardiac myofibroblasts from the Tabula Muris study (**Figures S8C-D**) while showing no appreciable expression in any cardiac populations from the Mouse Cell Atlas dataset (**Figures S8E-F**). We consider this evidence inconclusive in light of the discordant IHC patterns observed in HPA (not shown). This disagreement both within and across data types highlights the heart as a tissue which certainly requires thorough follow-up regarding the intricacies of ACE2 expression.

Across multiple studies of adipose tissue, ACE2 is not strongly expressed in any cell population (**Figure S9**). Of note, this includes studies of the murine adipose stromovascular fraction (**Figure S9A-D**) along with unfractionated adipose tissue subjected to single nucleus RNA-seq (sNuc-seq) which allows for the capture and sequencing of adipocytes (**Figure S9E-F**). DPP4 and ANPEP are both detected in adipose stromal populations along with smaller fractions of immune cells. Across multiple datasets, TMPRSS2 is exclusively expressed in cells defined by canonical epithelial markers (e.g. EPCAM, KRT8, CLDN3, KRT18), suggesting epithelial contamination of the adipose tissue preparations in these studies.

In the testis, ACE2 expression was unexpectedly low by scRNA-seq (**Figure S10A-D**) given its high expression by bulk RNA-seq (**Figure S3A**), strong staining by IHC (**Figure S10E)**, and high detection levels by proteomics (**Figures S3B-C**). The reason for this discrepancy is not clear. Based on the strong protein and bulk RNA-seq evidence, we suggest that SARS-CoV-2 may indeed be able to infect certain testicular cells, but our scRNA-seq analysis did not shed light on the most likely cellular targets in this case. In both human and mouse ovary, coronavirus entry receptors and TMPRSS2 are not appreciably expressed, which is consistent with a lack of detection by our other data modalities (**Figures S11A-D**).

In the liver, expression of these genes was generally consistent with the protein expression patterns observed by IHC (**Figure S12**). ACE2 shows minimal detection throughout liver populations, while both DPP4 and ANPEP are expressed in the epithelial compartment (**Figure S13**). ANPEP expression is particularly high in the $EPCAM^{+}$ population of cells which may mark hepatic progenitor cells. TMPRSS2 is also expressed in these cells and in a subset of mature hepatocytes, which is discordant with the lack of TMPRSS2 staining in the liver by IHC (**Figure S13**).

In the pancreas, ACE2 is expressed in a small fraction (~10%) of both acinar cells and ductal cells (**Figure S13**). DPP4 is robustly expressed in pancreatic alpha cells with lower expression detected in ~20% of ductal cells, while TMPRSS2 and ANPEP are both strongly expressed in the acinar and ductal populations (**Figure S13**). These patterns are largely consistent with our observations of protein expression by IHC (**Figure S12**).

To assess expression in blood and immune organs, we analyzed scRNA-seq studies from blood, spleen, bone marrow, and thymus **(Figures S14-15**). Generally ACE2 and TMPRSS2 are not highly expressed in any populations from these tissues. DPP4 is expressed in subsets of T cells across these studies (**Figure S15**) along with B cells and various progenitor populations in the bone marrow from the Tabula Muris study (**Figure S15**). ANPEP expression was mostly restricted to monocytes and macrophages along with some small bone marrow progenitor populations (**Figure S15B,D**). We conclude that immune cells are not likely targeted by SARS-CoV-2, but T cells could certainly serve as targets of MERS-CoV infection if the virus is able to enter circulation. Similarly, ANPEP expression in monocytes and macrophages may provide an alternative non-epithelial route of infection for other coronaviruses such as CoV-229E.

In both bladder and prostate samples from human and mouse, TMPRSS2 is robustly expressed in various epithelial populations (**Figures S16A-H**). The coronavirus entry receptors are only minimally detected across all bladder cell populations (**Figures S16A-F**), whereas DPP4 and ANPEP are detected in subsets of human prostate luminal cells (~8% of 18%, respectively) (**Figures S16H**). ACE2 expression is low across all recovered populations from the prostate (**Figure S16H**).

In the mouse uterus dataset of ~3,700 cells from the Mouse Cell Atlas (**Figure S17A**), Ace2 was uniformly absent from all recovered populations (**Figure S17B**). Anpep is robustly detected in ~60% of glandular epithelial cells along with a smaller fraction (~10%) of stromal cells (**Figure S17B**). Dpp4 expression is detected in ~17% of dendritic cells (n = 23 of 131) along with ~5% of glandular epithelial cells, and Tmprss2 is expressed in a similarly small fraction of the epithelial population (**Figure 17B**).

Finally we assessed the expression of these genes in central nervous system tissues despite their uniformly low expression and lack of detection in brain samples from GTEx and HPA, respectively (data not shown). scRNA-seq data suggests similarly low expression across CNS populations, including various regions of the mouse brain and the human retina (**Figure S18**). From the Tabula Muris study, Ace2 is detected in a small number of pericytes (22 of 146) and an even lower fraction of endothelial cells (**Figure S18B**). DPP4 is also expressed here in ~20% of endothelial cells (**Figure S18B**), which may warrant follow-up but importantly is not reflected in endothelial cells of the cerebral cortex by IHC (**Figure S19**). In all brain-derived cell populations from the Mouse Cell Atlas (**Figures S18C-D**) and human retina-derived populations (**Figures S18E-F**), these genes were uniformly not detected at meaningful levels.

**ACE2 expression and patient demographics**

While our study profiles the expression of the coronavirus receptors from various tissue samples, whether and how much receptor expression varies among individuals across various factors like age, disease states, genetic diversity, lifestyle, and environmental factors are not well understood. For instance, in order to understand variation of expression with age we explored ACE2 expression using GTEx samples. Interestingly, ACE2 expression levels in colon (transverse) samples were higher in younger individuals in comparison to older individuals (**Figure S24A**). In contrast, ACE2 expression levels in the esophagus (gastroesophageal junction) were lower in younger individuals in comparison to older individuals (**Figure S24B**). These patterns of ACE2 expression need to be tested more rigorously on larger sample sizes across diverse ethnicities to establish statistical significance. If ACE2 were found to be expressed more significantly in the colon of younger individuals, taken together with recent reports of sustained fecal shedding of SARS-CoV-2[10,11], the emerging epidemiological hypothesis as having lesser respiratory complications in general and serving as facile vectors in transmission of COVID-19 require more deeper investigation.

The recent reports of hypertension as a comorbidity in COVID-19 patients[12] and specifically the use of ACE1 inhibitors as antihypertensives contributing to mortality encourages a hypothesisfree examination of the FDA adverse event reporting system (FAERS; see below). Examining the differential patterns of adverse event reports between ACE1 inhibitors and beta blockers - both antihypertensive drug classes, with the former known to increase ACE2 expression in select cardiovascular tissues[13] - shows that ACE1 inhibitors have heightened respiratory oedema (**Figure S24C**). Any reports of patients on beta blockers also were weeded out of the ACE1 inhibitors set, and vice versa, for this analysis. The significantly amplified adverse events of epiglottic oedema, epiglottis, pneumopericardium, upper airway obstruction, eosinophilic oesophagitis, oedema mucosal, edema mouth, tracheal oedema, palatal oedema, and allergic oedema in ACE1 inhibitors compared to beta blockers suggests that a thorough investigation of all 10 million plus adverse event reports is necessary, to triangulate any drug-induced side effects that also appear as comorbidities from the emerging evidence of COVID-19 mortality. Availability of single-cell RNAseq data from healthy, pathological, and drug-treated tissues would enable us to profile the age-associated and treatment-based expression levels of ACE2 across cell populations. These observations clearly underline that urgent investments are needed to conduct comprehensive scRNAseq profiling of tissue samples from across different demographics and pathologies pertinent to COVID-19, as such effort will hold tremendous potential to reveal under-appreciated fingerprints of coronavirus transmission patterns, tissue tropism, and mortality.

#### FDA Adverse Event Reporting System (FAERS) analysis

The FAERS application of the nferX platform supports viewing adverse event profiles of all marketed products through multiple lenses - Count, Proportional Reporting Ratio (PRR), and an nferX Adverse Event (AE) Score. Count is the raw number of reports between a drug and an adverse event. The proportional reporting ratio (PRR) is a simple way to get a measure of how common an adverse event for a particular drug is compared to how common the event is in the overall database. A PRR > 1 for a drug-event combination indicates that a greater proportion of the reports for the drug are for the event than the proportion of events in the rest of the database, while a PRR of 2 for a drug event combination indicates that the proportion of reports for the drug-event combination is twice the proportion of the event in the overall database. The PRR is computed as follows:

m = number of reports with drug and event
n = number of reports with drug
M = number of reports with event in database
N = number of reports in database

Count of an event with a query drug is a good first measure of association. But it has the problem that generally common events will often show up at the top, where we're often more interested in events that are differentially associated with the query drug over other drugs. An issue with PRR is that it is noisy when the total number of event reports is small. If there are 3 reports of some oddly specific event and 1 occurs with the query drug, that event will likely have a very high PRR. But it may not be the event we'd be most interested in for a drug (in FAERS such rare events are often not even proper adverse events) - we want events that occur often, and also are differentially associated with a drug - a balance between count and PRR.

The AE score tries to strike this balance in an all-in-one measure. It up-weights events that occur often for the query drug (this is the ln(count) term), and that are differentially associated with the query drug (this is the sigmoid term).

The sigmoid(PRR-1.5) term ranges smoothly from 0 to 1. It's equal to 0.5 at PRR=1.5. When PRR=6, sigmoid(PRR-1.5)=0.99; so PRR values >= 6 are all treated roughly equivalently by the AE score. So extremely high PRRs due to small counts will not swing the AE score much beyond PRR=6. And the ln(count) term will down-weight those small-count cases, so that they do not show up at the top of the AE score list.

A nice property of AE score is that, for a given query drug, the AE scores of the events with that drug turn out to often roughly follow an exponential distribution, particularly at the tails. We can then fit exponential distributions to the scores, and look at them. A benefit of the exponential fit is that we can make more robust claims about how significant a certain score is for a query drug, even if the empirical data is sparse/noisy at the tails for a particular drug.

#### Expression of ACE2 in respiratory tract associated samples from GEO

In order to further assess this unexpected expression pattern, we also examined individual GEO studies with bulk RNA-seq data that are associated with the respiratory system (e.g. bronchi) using a keyword search. Specifically, we examined individual studies with bulk RNA-seq data from GEO that mentioned the following terms or their grammatical derivatives in the study title, study description, study design, or sample metadata: esophagus, alveoli, bronchial, lung, pulmonary, pharynx, oral, salivary, larynx, olfactory, trachea, nasal. This comprised nearly 200 unique studies in total.

While ACE2 expression was similarly low in the majority of these studies, we did find that ACE2 expression increased in bronchial epithelial cells over the course of differentiation in a cell culture system. However, this observation must be balanced against the large number of studies and samples that do *not* exhibit unexpectedly high expression levels, considering GTEx lung samples as the “baseline distribution.” More systematic studies of various respiratory tissues and cell types are warranted.

Among the studies with highest expression of ACE2 relative to the GTEx baseline, we call attention to the following (**Figure S25**):

- GSE113209 (n=96) includes nasal samples from healthy volunteers experimentally challenged with pneumococcus bacteria, three days after receiving live attenuated influenza vaccine or tetravalent inactivated influenza vaccine.
- GSE124949 (n=27) includes nasal samples from twenty adult healthy volunteers around the time of experimental human pneumococcal challenge. The samples were collected five days prior to, or two days after the experimental challenge.
- GSE97668 (n=33) includes nasal epithelial samples before and after asthmatics and non-asthmatic controls were inoculated with HRV-A16.
- GSE93526 (n=44) profiles the cells from air-liquid interface respiratory epithelial cell cultures which are infected with nontuberculous mycobacteria.

#### Supplementary Figures Legends

**Supplemental Figure 1. Validation of metrics used to assess literature-derived associations.** (A) Testing of cosine distance metric to capture global associations between two concepts. (B) Testing of exponential local score metric to capture proximity-based associations between concepts. (C) Comparing the histograms of partitioned cosine distance space for for a random set of word2vec vectors. (D) What does the word2vec neural network do? From the perspective of Genes-Diseases associations. (E) Studying how the number of dimensions being varied In the word2vec space modifies the cosine distance distributions for a large text corpus.

**Supplemental Figure 2. ACE2 expression in cell types from murine and human pancreas by scRNA-seq.** (A) Ace2 expression in pancreatic cell types from the Tabula Muris study[14], including polypeptide (gamma) and alpha cells. (B) ACE2 expression in pancreatic cell types from human samples[15–17], with notable absence in gamma and alpha cells.

**Supplemental Figure 3. Multimodal analysis of ACE2 expression using bulk RNA-seq, proteomics, and IHC.** (A) ACE2 expression by bulk RNA-seq across healthy tissues from GTEx. Gray box highlights the tissues with highest expression including small intestine and kidney; green arrow indicates lung. (B-C) Expression of ACE2 and other coronavirus receptors by proteomics across over 20 healthy human tissues from the Human Proteome Map (B) and the Human Protein Atlas (C). IHC stained images showing ACE2 expression in the kidney, small intestine, and colon, with minimal or now expression detected in respiratory tissues.

**Supplemental Figure 4. Overview of nferX Single Cell platform functionality.** Use case shows results for a query of *Ins1* in the Tabula Muris study. (**A**) Violin Plot section shows that pancreatic beta cells express *Ins1* at the highest level among 148 annotated cell types from 18 tissues. Summary statistics for each row indicate the percent of cells within a given cluster expressing *Ins1*, the mean expression of *Ins1* in that cluster, and a metric of expression specificity (Cohen's D) to each cluster. On the left side of the table, "Signals" columns highlight literature-derived associations between *Ins1* and the tissue ("pancreas") or cell type (e.g., "pancreatic beta cells"). (**B**) Dimensionality reduction-based visualization of all ~100,600 cells from Tabula Muris. Left-sided plot colors cells (points) based on *Ins1* expression in the individual cell; right-sided plot colors cells based on a selected metadata variable (here, tissue of origin). (**C**) List of cluster-defining genes for a selected cell population - by default, any cell population which highly expresses the query gene. Here, cluster-defining genes for pancreatic beta cells include *Ins1*, *Ins2*, and *Iapp*. Blue box highlights functionality to triangulate such cluster-defining gene lists with any literature query of interest; here, we see that *Ins1*, *Ins2*, *G6pc2*, and *Slc30a8* are not only specifically expressed in pancreatic beta cells but are also highly associated with the concept of "autoimmunity" throughout biomedical literature.

**Supplemental Figure 5. Assessment of DPP4, ANPEP, and TMPRSS2 across healthy tissues using bulk RNA-seq and IHC.** (A-C) GTEx tissues with highest expression observed for DPP4 (A), ANPEP (B), and TMPRSS2 (C). (D) Assessment of ACE2 expression along with other coronavirus receptors in non-GTEx tissues from the Janssen BodyMap dataset (bulk RNA-seq). (E) IHC staining showing that DPP4 and ANPEP are apically expressed in the small intestine (top), while DPP4, ANPEP, and TMPRSS2 are all expressed apically in renal tubular cells (bottom).

**Supplemental Figure 6. Single-cell RNAseq analysis of coronavirus receptors in the adult and fetal human kidney.** UMAP-based Dimensionality reduction plot denoting cell types (**A, G**) and corresponding expression levels of ACE2, DPP4, ANPEP and TMPRSS2 (**B, H**) from adult (**A-B**) and fetal (**G-H**) human kidney samples[9]. (**C-F**) Violin plot visualization of gene expression for ACE2 (**C**), TMPRSS2 (**D**), ANPEP (**E**), and DPP4 (**F**) in adult human kidney scRNA-seq dataset. Automated literature synthesis highlights strong associations of ACE2, DPP4, and ANPEP to the kidney.

**Supplemental Figure 7. ACE2, DPP4, and ANPEP show similar expression profiles in the renal proximal tubule epithelial cells by scRNA-seq and bulk RNA-seq.** (**A-C**) Cosine similarities between gene expression vectors of all genes and ACE2 (**A**), DPP4 (**B**), and ANPEP (**C**) among all annotated proximal tubule epithelial cells from a human kidney scRNA-seq dataset[9]. ACE2, DPP4, and ANPEP are reciprocally among the most similar genes to each other. (**D-F**) Analysis of pearson correlation coefficients between all genes and ACE2 (**G**), DPP4 (**H**), and ANPEP (**I**) from human kidney cortex samples (GTEx; n = 85). ACE2, DPP4, and ANPEP are reciprocally among the most strongly correlated genes to each other.

**Supplemental Figure 8. Single-cell RNAseq analysis of coronavirus receptors in the human and murine heart.** UMAP-based Dimensionality reduction plot denoting cell types (**A, C, E**) and corresponding expression levels of ACE2, DPP4, ANPEP and TMPRSS2 (**B, D, F**) from multiple studies: (**A-B**) human heart[18], (**C-D**) murine heart from Tabula Muris[14], and (E-F) murine heart from Mouse Cell Atlas[19].

**Supplemental Figure 9. Single-cell RNAseq analysis of coronavirus receptors in adipose tissue.** UMAP-based Dimensionality reduction plot denoting cell types (**A, C, E**) and corresponding expression levels of Ace2, Dpp4, Anpep and Tmprss2 (**B, D, F**) from multiple studies: (**A-B**) adipose stromovascular fraction from mice treated with PBS or a beta-3 adrenergic agonist[20], (**C-D**) murine adipose stromovascular fraction from Tabula Muris[14], and (**E-F**) adipose tissue from mice subjected to 24 hours of cold shock and processed for single nucleus RNA-sequencing[20]. (**G-H**) Identification of adipocytes as the most likely ACE2-expressing cell type in adipose tissue based on literature associations. The query shown is from the nferX Signals platform and effectively extracts any textual fragments which contain [“ACE2” or its gene synonyms] *AND* [displayed terms related to gene expression/protein detection] *AND* [“adipose” *OR* “fat]. Among these textual fragments, we extract the tokens which represent cell types and calculate enrichments of each cell type among these fragments, normalized by the number of occurrences of this cell type elsewhere in the literature.

**Supplemental Figure 10. Single-cell RNAseq analysis of coronavirus receptors in the human and murine testis.** UMAP-based Dimensionality reduction plot denoting cell types (**A, C**) and corresponding expression levels of ACE2, DPP4, ANPEP and TMPRSS2 (**B, D**) from multiple studies: (**A-B**) adult human testis[21], and (**C-D**) murine testis from Mouse Cell Atlas[19].

**Supplemental Figure 11. Single-cell RNAseq analysis of coronavirus receptors in the human and murine ovary.** UMAP-based Dimensionality reduction plot denoting cell types (**A, C**) and corresponding expression levels of ACE2, DPP4, ANPEP and TMPRSS2 (**B, D**) from multiple studies: (**A-B**) adult human ovary[22], and (**C-D**) murine ovary from Mouse Cell Atlas[19].

**Supplemental Figure 12. IHC images of coronavirus receptors in healthy pancreas and liver samples from the Human Protein Atlas.** (A-B) ACE2 expression on the apical surface of gallbladder epithelial cells (A) and on ductal surfaces in the pancreas (B). (C) ANPEP is expressed on apical membranes in pancreatic acini and ducts. (D) TMPRSS2 is expressed on apical membranes in pancreatic acini and ducts. (E) DPP4 is expressed weakly in the pancreas, both on ductal surfaces and in endocrine islets. (F) DPP4 expression in healthy liver. (G) ANPEP expression in healthy liver; ANPEP appears to be the most strongly expressed coronavirus receptor in the human liver.

**Supplemental Figure 13. Single-cell RNAseq analysis of coronavirus receptors in the human liver and pancreas.** UMAP-based Dimensionality reduction plot denoting cell types (**A, C**) and corresponding expression levels of ACE2, DPP4, ANPEP and TMPRSS2 (**B, D**) from multiple studies: (**A-B**) human liver[23], and (**C-D**) three integrated studies of human pancreas[15–17].

**Supplemental Figure 14. Single-cell RNAseq analysis of coronavirus receptors in human blood, spleen, and bone marrow.** UMAP-based Dimensionality reduction plot denoting cell types (**A, C, E**) and corresponding expression levels of ACE2, DPP4, ANPEP and TMPRSS2 (**B, D, F**) from multiple studies: (**A-B**) human peripheral blood mononuclear cells[24], (**C-D**) human spleen[25], and (**E-F**) human bone marrow[26].

**Supplemental Figure 15. Single-cell RNAseq analysis of coronavirus receptors in murine spleen, bone marrow, and thymus.** UMAP-based Dimensionality reduction plot denoting cell types (**A, C, E**) and corresponding expression levels of ACE2, DPP4, ANPEP and TMPRSS2 (**B, D, F**) from multiple tissues in the Tabula Muris study[14]: (**A-B**) spleen, (**C-D**) bone marrow, (**E-F**) thymus.

**Supplemental Figure 16. Single-cell RNAseq analysis of coronavirus receptors in human and murine bladder and prostate.** UMAP-based Dimensionality reduction plot denoting cell types (**A, C, E, G**) and corresponding expression levels of ACE2, DPP4, ANPEP and TMPRSS2 (**B, D, F, H**) from multiple studies: (**A-B**) human bladder[27], (**C-D**) murine bladder from Tabula Muris[14], (**E-F**) murine bladder from Mouse Cell Atlas[19], and (**G-H**) human prostate and prostatic urethra.

**Supplemental Figure 17. Single-cell RNAseq analysis of coronavirus receptors in the murine uterus.** UMAP-based Dimensionality reduction plot denoting cell types (**A**) and corresponding expression levels of ACE2, DPP4, ANPEP and TMPRSS2 (**B**) in uterus-derived cells from the Mouse Cell Atlas[19].

**Supplemental Figure 18. Single-cell RNAseq analysis of coronavirus receptors in human and murine central nervous system tissues.** UMAP-based Dimensionality reduction plot denoting cell types (**A, C, E**) and corresponding expression levels of ACE2, DPP4, ANPEP and TMPRSS2 (**B, D, F**) in CNS samples from multiple studies: (A-B) murine brain from Tabula Muris[14], (C-D) murine brain from Mouse Cell Atlas[19], and (E-F) human retina[28].

**Supplemental Figure 19. IHC analysis of DPP4 expression in human brain cortex.** Images from the Human Protein Atlas showing brain cortex samples from three different individuals stained to assess DPP4 expression. Staining is notably absent from all cell types in all samples tested.

**Supplemental Figure 20. ACE2 is strongly correlated to surfactant protein-encoding genes across GTEx lung samples.** Distribution of pearson correlation coefficients between expression (in TPM) of ACE2 and all other genes detected with a mean of at least 1 TPM in the GTEx lung study (n = 573). The right tail of the distribution (shaded red) includes the top 4% of correlated genes (top). Each gene encoding a known surfactant protein is strongly correlated to ACE2 in lung samples, with the pearson correlation coefficient among this top 4%.

**Supplemental Figure 21. Negative IHC staining of ACE2 in nasopharynx from Human Protein Atlas.** Shown are two samples from different patients stained with different anti-ACE2 antibodies.

**Supplemental Figure 22. Coronavirus receptors show highly correlated expression patterns by single cell and bulk RNA-seq in human small intestine.** (**A**) Distribution of cosine distances between the 'gene expression vectors' of ACE2 and all genes in a scRNA-seq study of the human small intestine. (**B**) Genes similar to ACE2 sorted by literature-derivation association. (**C**) Gene expression correlations to ACE2 from bulk RNA-sequencing (GTEx) of human small intestine samples.

.

**Supplemental Figure 23. Characterization of oral epithelium cluster-defining genes from Hao, et al., 2020[29].** From a non-public scRNA-seq study, oral mucosal epithelial cells (including tongue-derived epithelial cells) were defined by expression of SFN, KRT6A, and KRT10. Assessment of SFN (A), KRT6A (B), and KRT10 (C) expression in tongue cells sequenced in the Tabula Muris study[14]. Each gene is most strongly expressed in murine tongue keratinocytes, with expression in basal epidermal cells as well.

**Supplemental Figure 24. Association of age and co-administered drugs with ACE2 expression of COVID-19 outcomes.** (**A**) Among GTEx transverse colon samples (n = 405), ACE2 expression tends to be higher in samples derived from younger individuals. (**B**) Among GTEx gastroesophageal junction samples, ACE2 expression tends to be higher in samples derived from older individuals. (**C**) Analysis of differential adverse events between ACE inhibitors and other classes of antihypertensive medications (example shown for beta-blockers), real-world evidence shows that patients taking ACE inhibitors are more likely to experience angioedema of various tissues including the small intestine and epiglottis.

**Supplemental Figure 25. Expression of ACE2 in respiratory tract associated samples from GEO in comparison Lung (GTEx).** Plots showing the comparison expression distribution of ACE2 between samples from lung (GTEx in purple) and samples from selected GEO studies (in green). The GEO identifiers for each comparison is shown.

#### References

1. Park, J. *et al.* Recapitulation and Retrospective Prediction of Biomedical Associations Using Temporally-enabled Word Embeddings. doi:10.1101/627513.
2. Mikolov, T., Sutskever, I., Chen, K., Corrado, G. & Dean, J. Distributed Representations of Words and Phrases and their Compositionality. (2013).
3. Cohen's D: Definition, Examples, Formulas - Statistics How To. *Statistics How To* https://www.statisticshowto.datasciencecentral.com/cohens-d/ (2016).
4. Contributors to Wikimedia projects. Mann–Whitney U test - Wikipedia. *Wikimedia Foundation, Inc.* https://en.wikipedia.org/wiki/Mann%E2%80%93Whitney_U_test (2004).
5. Contributors to Wikimedia projects. Brier score - Wikipedia. *Wikimedia Foundation, Inc.* https://en.wikipedia.org/wiki/Brier_score (2005).
6. dansbecker. What is Log Loss? *Kaggle* https://kaggle.com/dansbecker/what-is-log-loss (2018).
7. St John, J., Powell, K., Conley-Lacomb, M. K. & Chinni, S. R. TMPRSS2-ERG Fusion Gene Expression in Prostate Tumor Cells and Its Clinical and Biological Significance in Prostate Cancer Progression. *J. Cancer Sci. Ther.* **4**, 94–101 (2012).
8. Tabula Muris Consortium *et al.* Single-cell transcriptomics of 20 mouse organs creates a Tabula Muris. *Nature* **562**, 367–372 (2018).
9. Stewart BJ, E. al. Spatiotemporal immune zonation of the human kidney. - PubMed - NCBI. https://www.ncbi.nlm.nih.gov/pubmed/?term=31604275.
10. Xu, Y. *et al.* Characteristics of pediatric SARS-CoV-2 infection and potential evidence for persistent fecal viral shedding. *Nature Medicine* (2020) doi:10.1038/s41591-020-0817-4.
11. Gu, J., Han, B. & Wang, J. COVID-19: Gastrointestinal manifestations and potential fecal-oral transmission. *Gastroenterology* (2020) doi:10.1053/j.gastro.2020.02.054.
12. Fang, L., Karakiulakis, G. & Roth, M. Are patients with hypertension and diabetes mellitus at increased risk for COVID-19 infection? *The Lancet Respiratory Medicine* (2020) doi:10.1016/s2213-2600(20)30116-8.
13. Ferrario, C. M. *et al.* Effect of angiotensin-converting enzyme inhibition and angiotensin II receptor blockers on cardiac angiotensin-converting enzyme 2. *Circulation* **111**, 2605–2610 (2005).
14. Tabula Muris Consortium , et al. Single-cell transcriptomics of 20 mouse organs creates a Tabula Muris. - PubMed - NCBI. https://www.ncbi.nlm.nih.gov/pubmed/30283141.
15. Muraro MJ, E. al. A Single-Cell Transcriptome Atlas of the Human Pancreas. - PubMed - NCBI. https://www.ncbi.nlm.nih.gov/pubmed/?term=27693023.
16. Segerstolpe Å, E. al. Single-Cell Transcriptome Profiling of Human Pancreatic Islets in Health and Type 2 Diabetes. - PubMed - NCBI. https://www.ncbi.nlm.nih.gov/pubmed/?term=27667667.
17. Grün D, E. al. De Novo Prediction of Stem Cell Identity using Single-Cell Transcriptome Data. - PubMed - NCBI. https://www.ncbi.nlm.nih.gov/pubmed/?term=27345837.
18. Wang L, E. al. Single-cell reconstruction of the adult human heart during heart failure and recovery reveals the cellular landscape underlying cardiac function. - PubMed - NCBI. https://www.ncbi.nlm.nih.gov/pubmed/?term=31915373.
19. Han X, E. al. Mapping the Mouse Cell Atlas by Microwell-Seq. - PubMed - NCBI. https://www.ncbi.nlm.nih.gov/pubmed/29474909.
20. Rajbhandari P, E. al. Single cell analysis reveals immune cell-adipocyte crosstalk regulating the transcription of thermogenic adipocytes. - PubMed - NCBI. https://www.ncbi.nlm.nih.gov/pubmed/?term=31644425.

21. Guo J, E. al. The adult human testis transcriptional cell atlas. - PubMed - NCBI. https://www.ncbi.nlm.nih.gov/pubmed/?term=30315278.
22. Fan X, E. al. Single-cell reconstruction of follicular remodeling in the human adult ovary. - PubMed - NCBI. https://www.ncbi.nlm.nih.gov/pubmed/?term=31320652.
23. Aizarani N, E. al. A human liver cell atlas reveals heterogeneity and epithelial progenitors. - PubMed - NCBI. https://www.ncbi.nlm.nih.gov/pubmed/?term=31292543.
24. Single Cell Portal. https://singlecell.broadinstitute.org/single_cell/study/SCP345/ica-blood-mononuclear-cells-2-donors-2-sites.
25. Website. https://www.biorxiv.org/content/10.1101/741405v1.
26. HCA Data Browser. https://data.humancellatlas.org/explore/projects/cc95ff89-2e68-4a08-a234-480eca21ce79.
27. Yu Z, E. al. Single-Cell Transcriptomic Map of the Human and Mouse Bladders. - PubMed - NCBI. https://www.ncbi.nlm.nih.gov/pubmed/31462402.
28. Menon M, E. al. Single-cell transcriptomic atlas of the human retina identifies cell types associated with age-related macular degeneration. - PubMed - NCBI. https://www.ncbi.nlm.nih.gov/pubmed/?term=31653841.
29. Xu, H. *et al.* High expression of ACE2 receptor of 2019-nCoV on the epithelial cells of oral mucosa. *Int. J. Oral Sci.* **12**, 1–5 (2020).

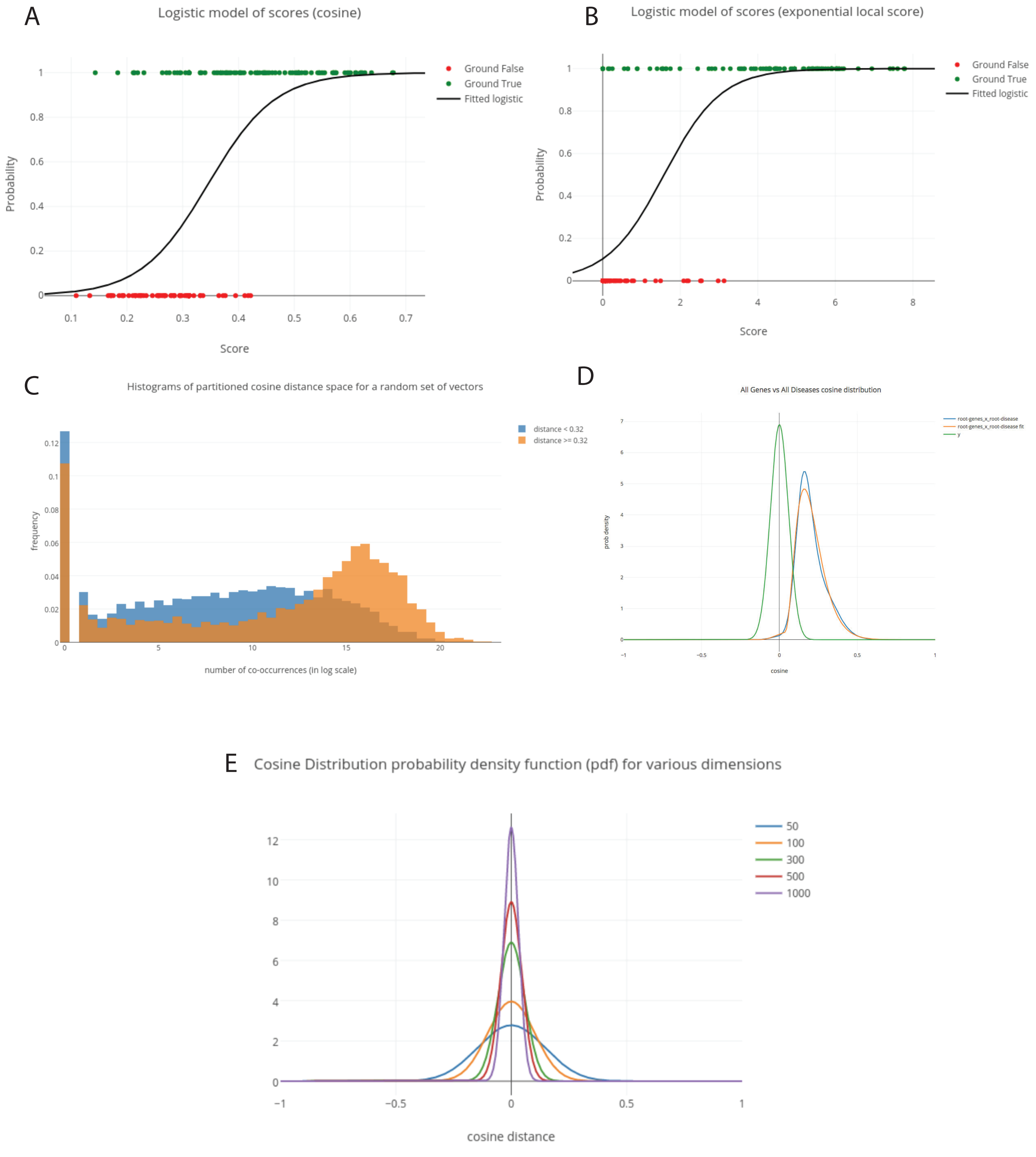


Supplemental Figure 1

A

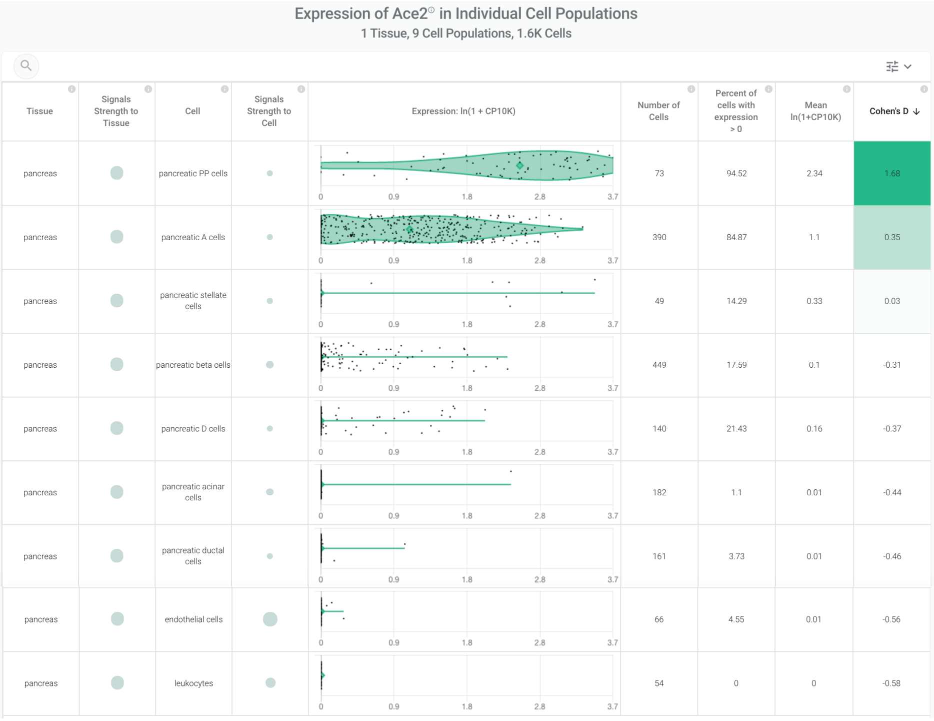

Expression of Ace2 in Individual Cell Populations
1 Tissue, 9 Cell Populations, 1.6K Cells

| Tissue | Signals Strength to Tissue | Cell | Signals Strength to Cell | Expression: ln(1 + CP10K) | Number of Cells | Percent of cells with expression > 0 | Mean ln(1+CP10K) | Cohen's D ↓ |
|---|---|---|---|---|---|---|---|---|
| pancreas | | pancreatic PP cells | | | 73 | 94.52 | 2.34 | 1.68 |
| pancreas | | pancreatic A cells | | | 390 | 84.87 | 1.1 | 0.35 |
| pancreas | | pancreatic stellate cells | | | 49 | 14.29 | 0.33 | 0.03 |
| pancreas | | pancreatic beta cells | | | 449 | 17.59 | 0.1 | -0.31 |
| pancreas | | pancreatic D cells | | | 140 | 21.43 | 0.16 | -0.37 |
| pancreas | | pancreatic acinar cells | | | 182 | 1.1 | 0.01 | -0.44 |
| pancreas | | pancreatic ductal cells | | | 161 | 3.73 | 0.01 | -0.46 |
| pancreas | | endothelial cells | | | 66 | 4.55 | 0.01 | -0.56 |
| pancreas | | leukocytes | | | 54 | 0 | 0 | -0.58 |

B

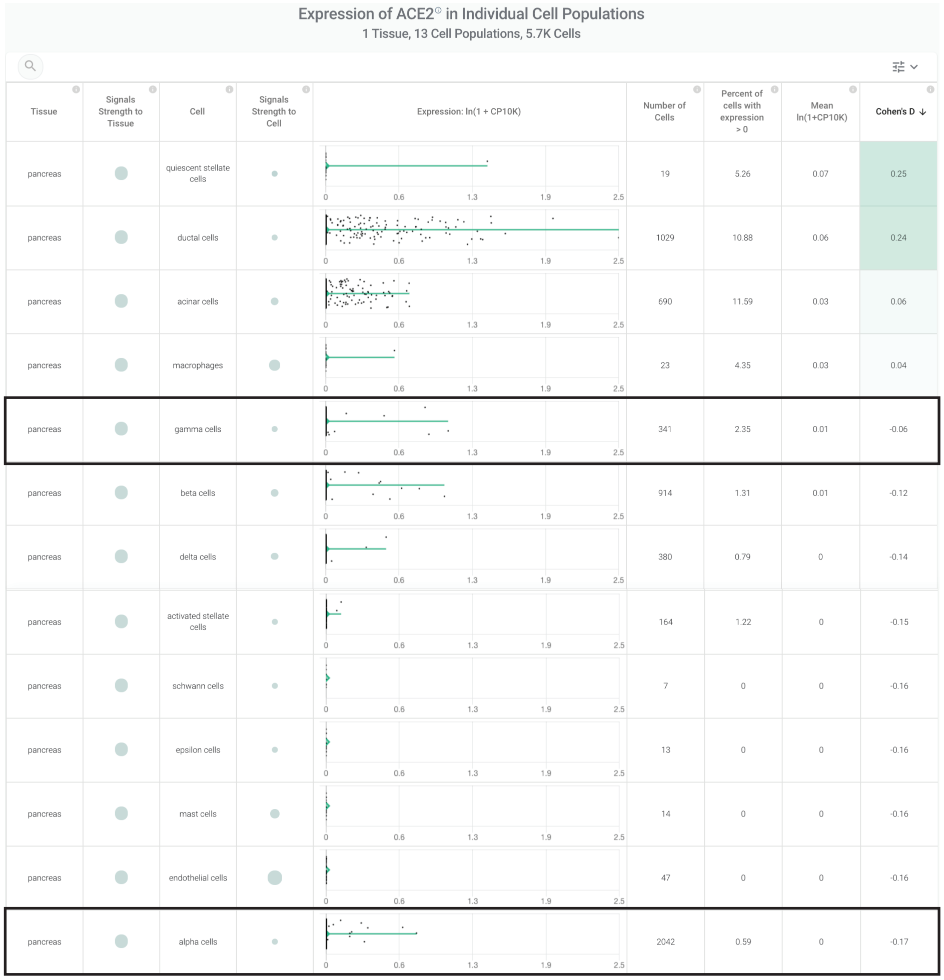

Expression of ACE2 in Individual Cell Populations
1 Tissue, 13 Cell Populations, 5.7K Cells

| Tissue | Signals Strength to Tissue | Cell | Signals Strength to Cell | Expression: ln(1 + CP10K) | Number of Cells | Percent of cells with expression > 0 | Mean ln(1+CP10K) | Cohen's D ↓ |
|---|---|---|---|---|---|---|---|---|
| pancreas | | quiescent stellate cells | | | 19 | 5.26 | 0.07 | 0.25 |
| pancreas | | ductal cells | | | 1029 | 10.88 | 0.06 | 0.24 |
| pancreas | | acinar cells | | | 690 | 11.59 | 0.03 | 0.06 |
| pancreas | | macrophages | | | 23 | 4.35 | 0.03 | 0.04 |
| pancreas | | gamma cells | | | 341 | 2.35 | 0.01 | -0.06 |
| pancreas | | beta cells | | | 914 | 1.31 | 0.01 | -0.12 |
| pancreas | | delta cells | | | 380 | 0.79 | 0 | -0.14 |
| pancreas | | activated stellate cells | | | 164 | 1.22 | 0 | -0.15 |
| pancreas | | schwann cells | | | 7 | 0 | 0 | -0.16 |
| pancreas | | epsilon cells | | | 13 | 0 | 0 | -0.16 |
| pancreas | | mast cells | | | 14 | 0 | 0 | -0.16 |
| pancreas | | endothelial cells | | | 47 | 0 | 0 | -0.16 |
| pancreas | | alpha cells | | | 2042 | 0.59 | 0 | -0.17 |

Supplemental Figure 2

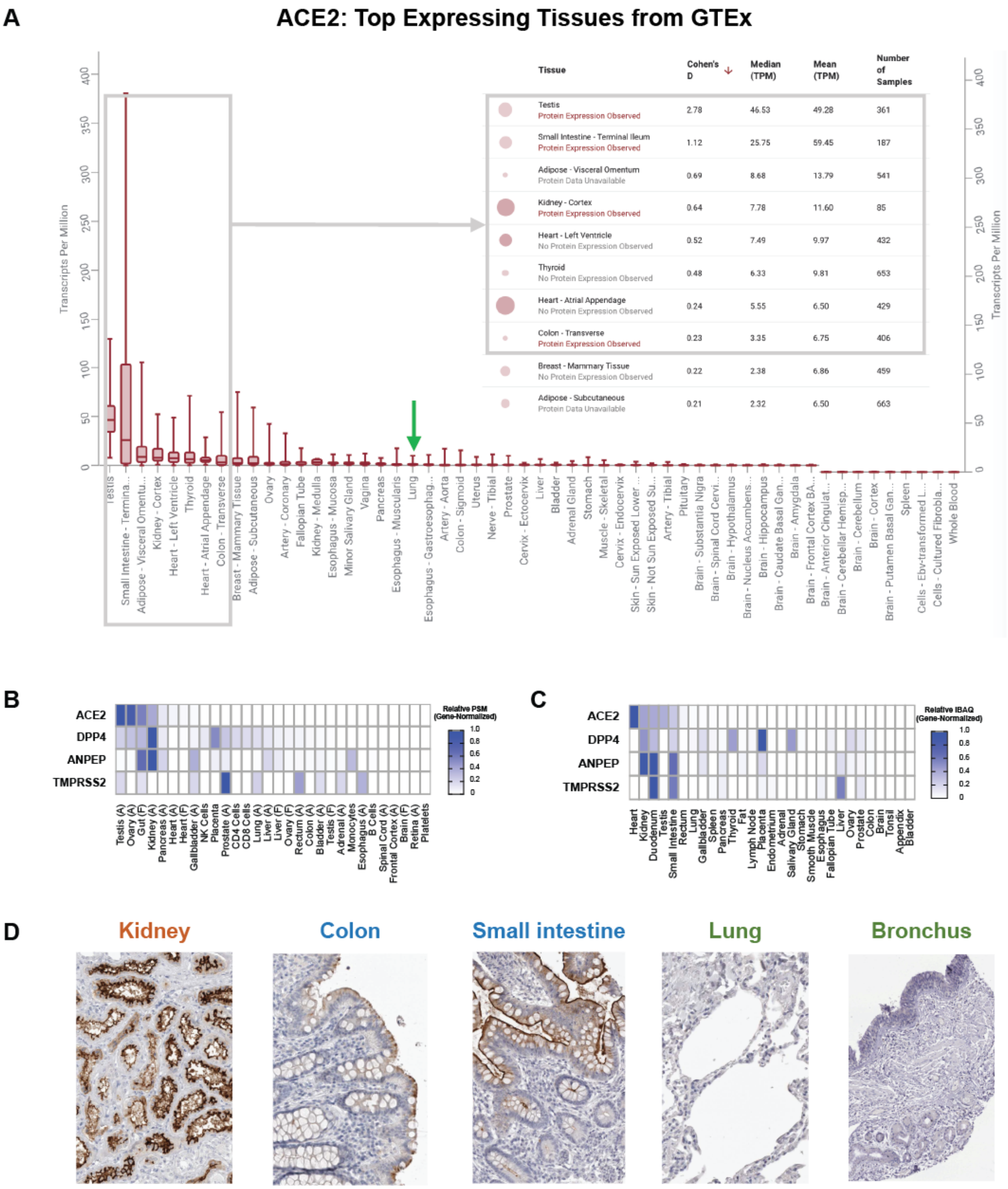


Supplemental Figure 3

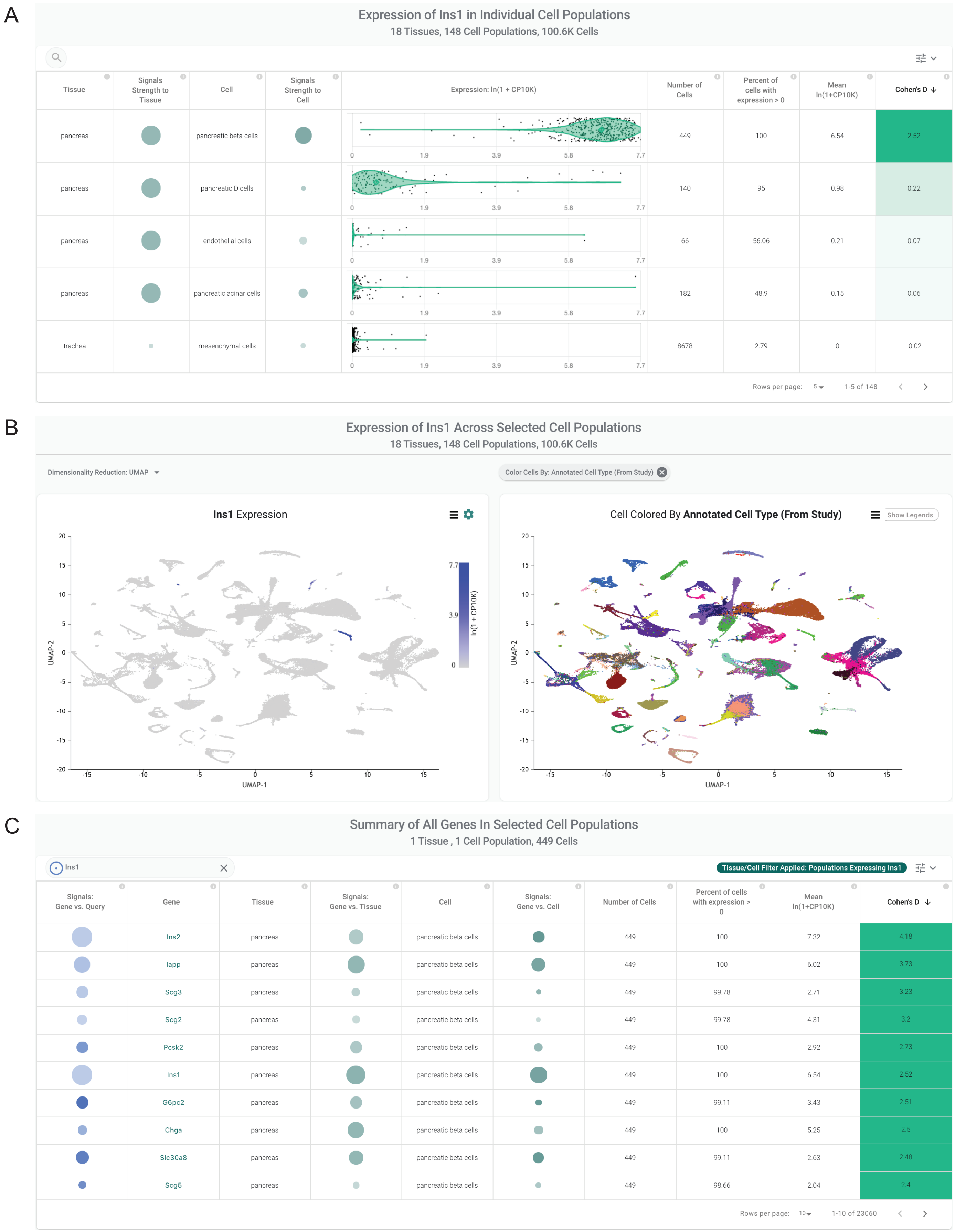


| Tissue | Signals Strength to Tissue | Cell | Signals Strength to Cell | Expression: ln(1 + CP10K) | Number of Cells | Percent of cells with expression > 0 | Mean ln(1+CP10K) | Cohen's D |
|---|---|---|---|---|---|---|---|---|
| pancreas | | pancreatic beta cells | | | 449 | 100 | 6.54 | 2.52 |
| pancreas | | pancreatic D cells | | | 140 | 95 | 0.98 | 0.22 |
| pancreas | | endothelial cells | | | 66 | 56.06 | 0.21 | 0.07 |
| pancreas | | pancreatic acinar cells | | | 182 | 48.9 | 0.15 | 0.06 |
| trachea | | mesenchymal cells | | | 8678 | 2.79 | 0 | -0.02 |



| Signals: Gene vs. Query | Gene | Tissue | Signals: Gene vs. Tissue | Cell | Signals: Gene vs. Cell | Number of Cells | Percent of cells with expression > 0 | Mean ln(1+CP10K) | Cohen's D |
|---|---|---|---|---|---|---|---|---|---|
| | Ins2 | pancreas | | pancreatic beta cells | | 449 | 100 | 7.32 | 4.18 |
| | Iapp | pancreas | | pancreatic beta cells | | 449 | 100 | 6.02 | 3.73 |
| | Scg3 | pancreas | | pancreatic beta cells | | 449 | 99.78 | 2.71 | 3.23 |
| | Scg2 | pancreas | | pancreatic beta cells | | 449 | 99.78 | 4.31 | 3.2 |
| | Pcsk2 | pancreas | | pancreatic beta cells | | 449 | 100 | 2.92 | 2.73 |
| | Ins1 | pancreas | | pancreatic beta cells | | 449 | 100 | 6.54 | 2.52 |
| | G6pc2 | pancreas | | pancreatic beta cells | | 449 | 99.11 | 3.43 | 2.51 |
| | Chga | pancreas | | pancreatic beta cells | | 449 | 100 | 5.25 | 2.5 |
| | Slc30a8 | pancreas | | pancreatic beta cells | | 449 | 99.11 | 2.63 | 2.48 |
| | Scg5 | pancreas | | pancreatic beta cells | | 449 | 98.66 | 2.04 | 2.4 |



Supplemental Figure 4

A

#### DPP4: Top Expressing Tissues from GTEx

| Tissue | Cohen's D ↓ | Median (TPM) | Mean (TPM) | Number of Samples |
|---|---|---|---|---|
| Cells - Cultured Fibroblasts<br>Protein Data Unavailable | 1.44 | 48.83 | 64.12 | 504 |
| Small Intestine - Terminal Ileum<br>Protein Expression Observed | 1.10 | 28.24 | 48.68 | 187 |
| Prostate<br>Protein Expression Observed | 1.04 | 28.25 | 42.06 | 245 |
| Kidney - Cortex<br>Protein Expression Observed | 0.90 | 19.71 | 28.92 | 85 |
| Adipose - Subcutaneous<br>Protein Data Unavailable | 0.79 | 16.67 | 25.15 | 663 |
| Adipose - Visceral Omentum<br>Protein Data Unavailable | 0.58 | 12.55 | 17.96 | 541 |
| Lung<br>No Protein Expression Observed | 0.37 | 10.73 | 13.65 | 578 |

B

#### ANPEP: Top Expressing Tissues from GTEx

| Tissue | Cohen's D ↓ | Median (TPM) | Mean (TPM) | Number of Samples |
|---|---|---|---|---|
| Cells - Cultured Fibroblasts<br>Protein Data Unavailable | 3.21 | 810.50 | 874.09 | 504 |
| Pancreas<br>Protein Expression Observed | 1.67 | 386.05 | 414.82 | 328 |
| Small Intestine - Terminal Ileum<br>Protein Expression Observed | 1.19 | 634.00 | 1,047.39 | 187 |
| Kidney - Cortex<br>Protein Expression Observed | 0.68 | 176.30 | 215.45 | 85 |
| Whole Blood<br>Protein Data Unavailable | 0.56 | 159.00 | 179.33 | 755 |
| Liver<br>Protein Expression Observed | 0.55 | 159.80 | 173.82 | 226 |
| Colon - Transverse<br>Protein Expression Observed | 0.27 | 64.96 | 133.32 | 406 |

C

#### TMPRSS2: Top Expressing Tissues from GTEx

| Tissue | Cohen's D ↓ | Median (TPM) | Mean (TPM) | Number of Samples |
|---|---|---|---|---|
| Prostate<br>Protein Expression Observed | 1.44 | 178.10 | 228.82 | 245 |
| Stomach<br>Protein Expression Observed | 1.43 | 113.60 | 108.97 | 359 |
| Colon - Transverse<br>No Protein Expression Observed | 0.94 | 63.85 | 83.59 | 406 |
| Pancreas<br>Protein Expression Observed | 0.94 | 48.68 | 49.76 | 328 |
| Lung<br>No Protein Expression Observed | 0.86 | 43.22 | 49.04 | 578 |
| Small Intestine - Terminal Ileum<br>Protein Expression Observed | 0.85 | 42.10 | 63.19 | 187 |
| Minor Salivary Gland<br>Protein Expression Observed | 0.71 | 41.44 | 40.07 | 162 |
| Kidney - Medulla<br>Protein Expression Observed | 0.61 | 30.04 | 37.44 | 4 |

D

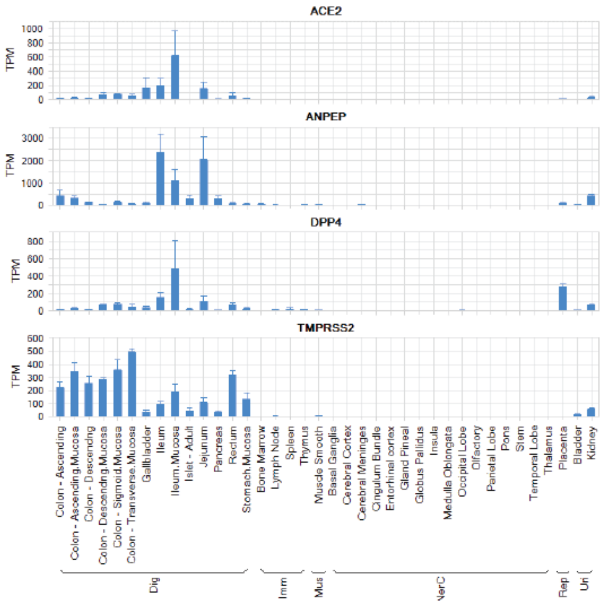


E

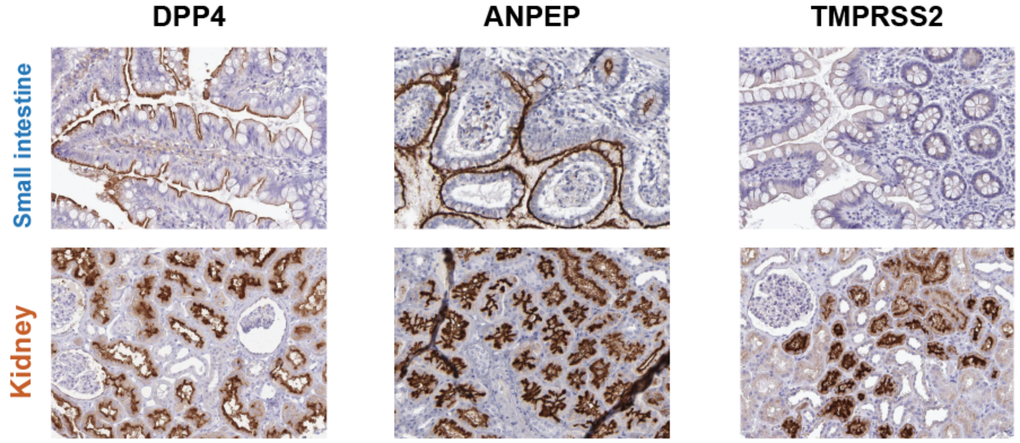


Supplemental Figure 5

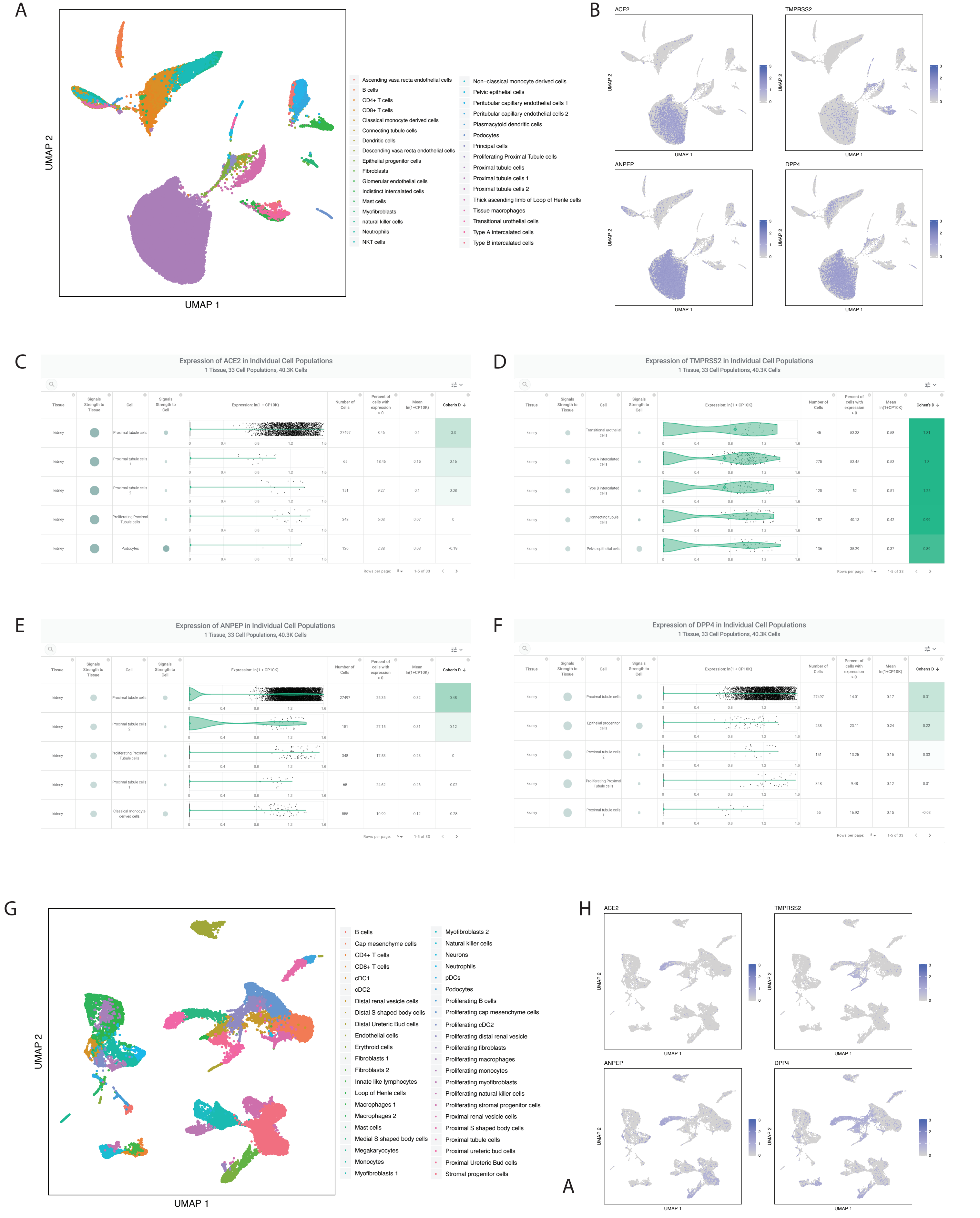


Supplemental Figure 6

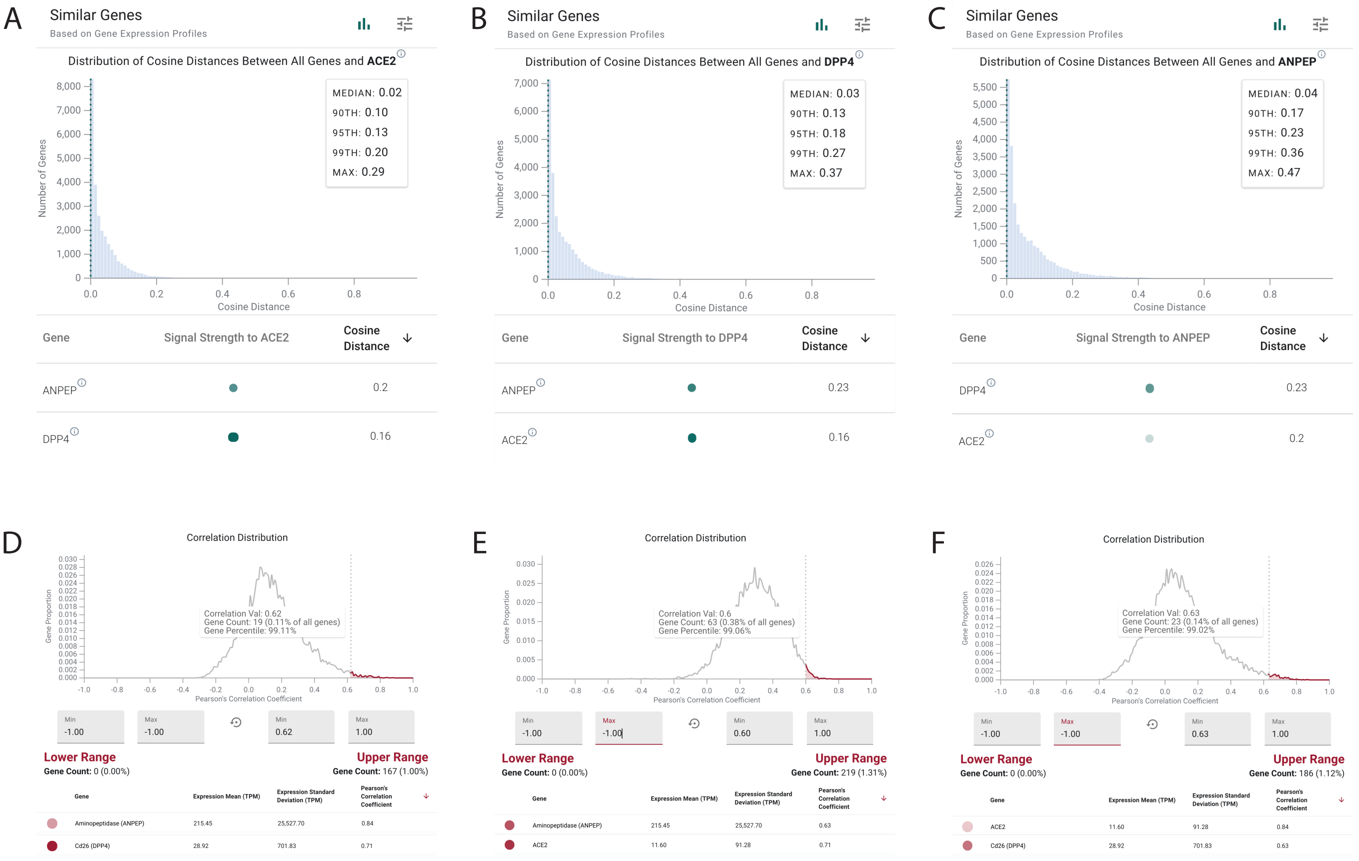


Supplemental Figure 7

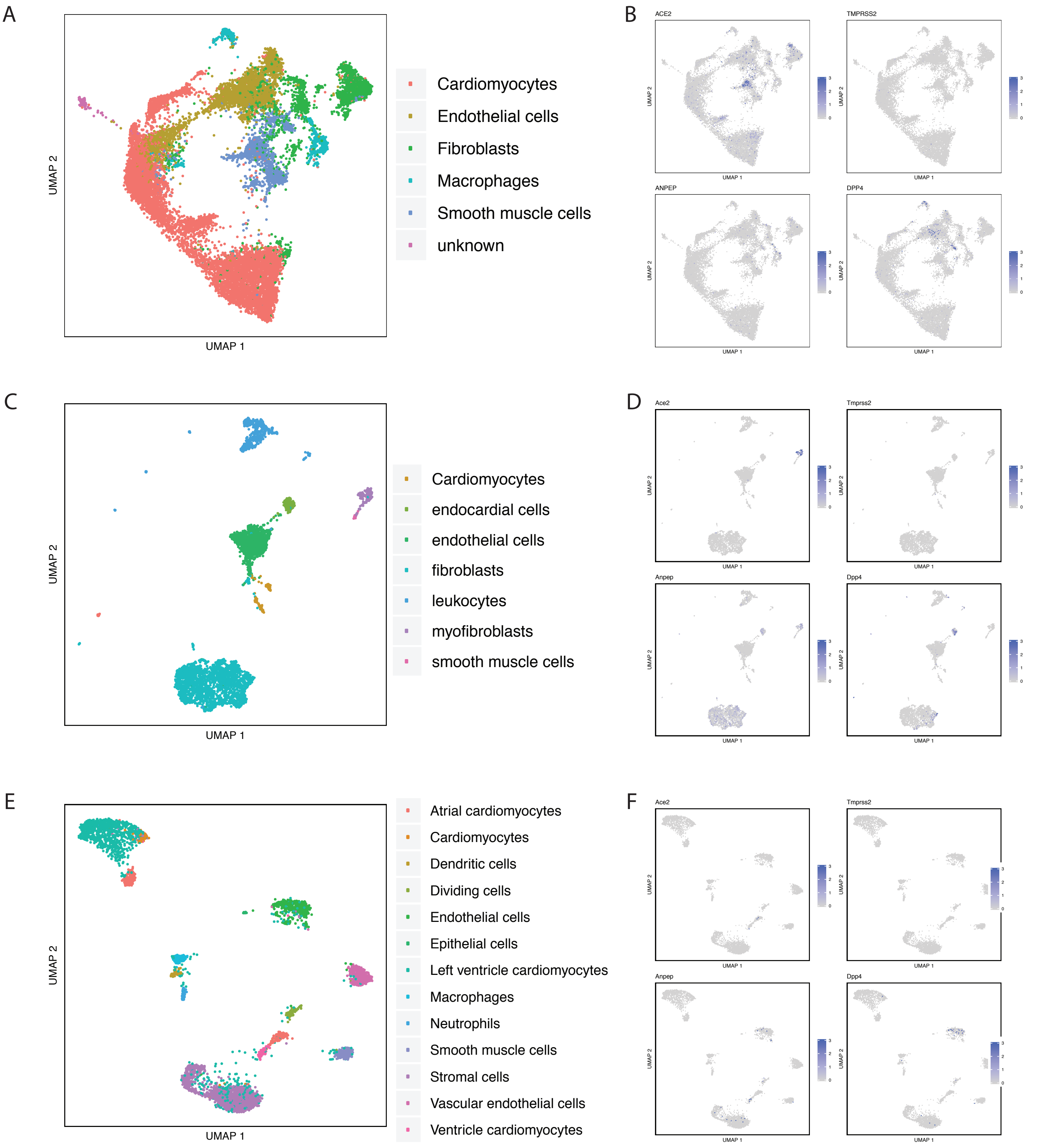


Supplemental Figure 8

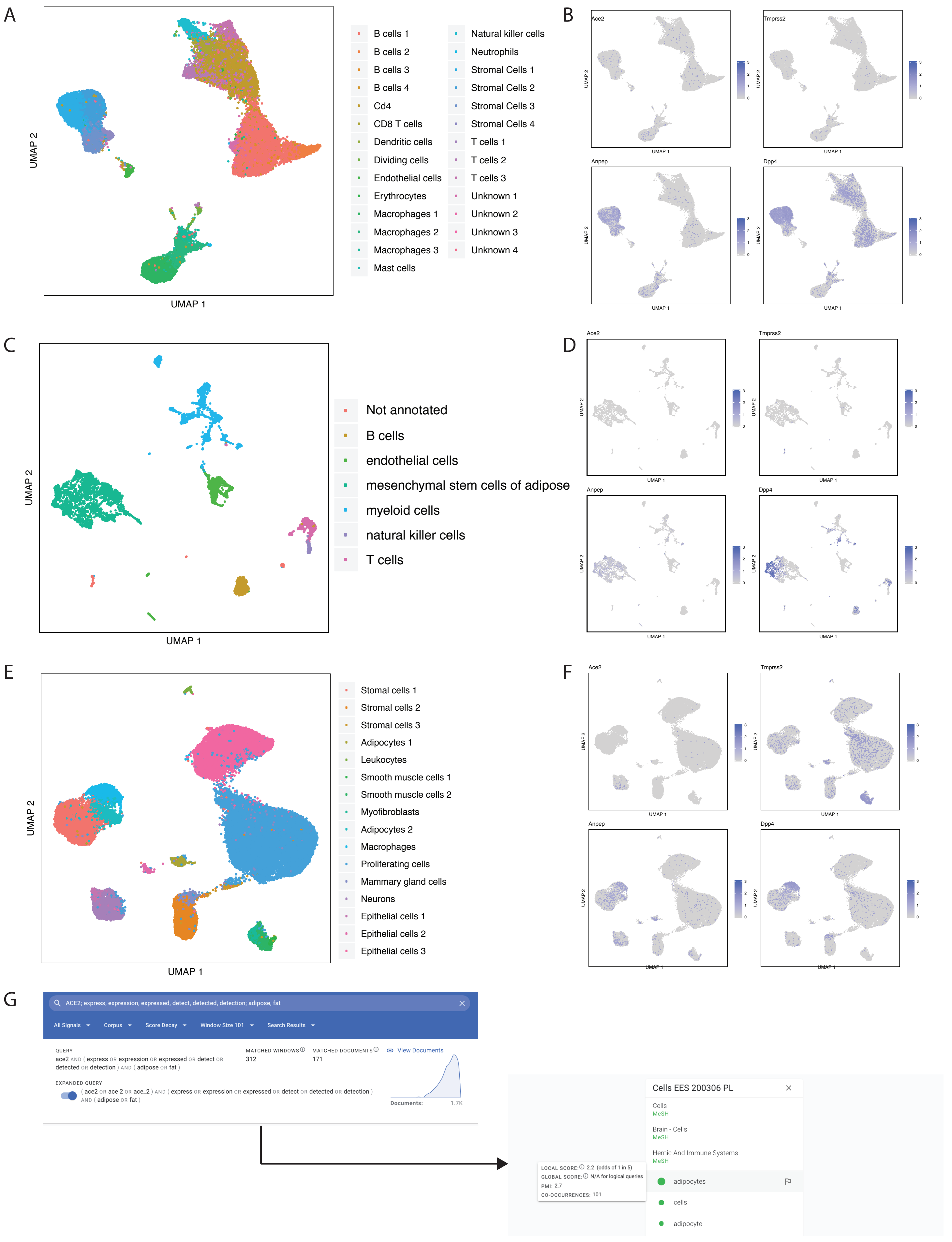


Supplemental Figure 9

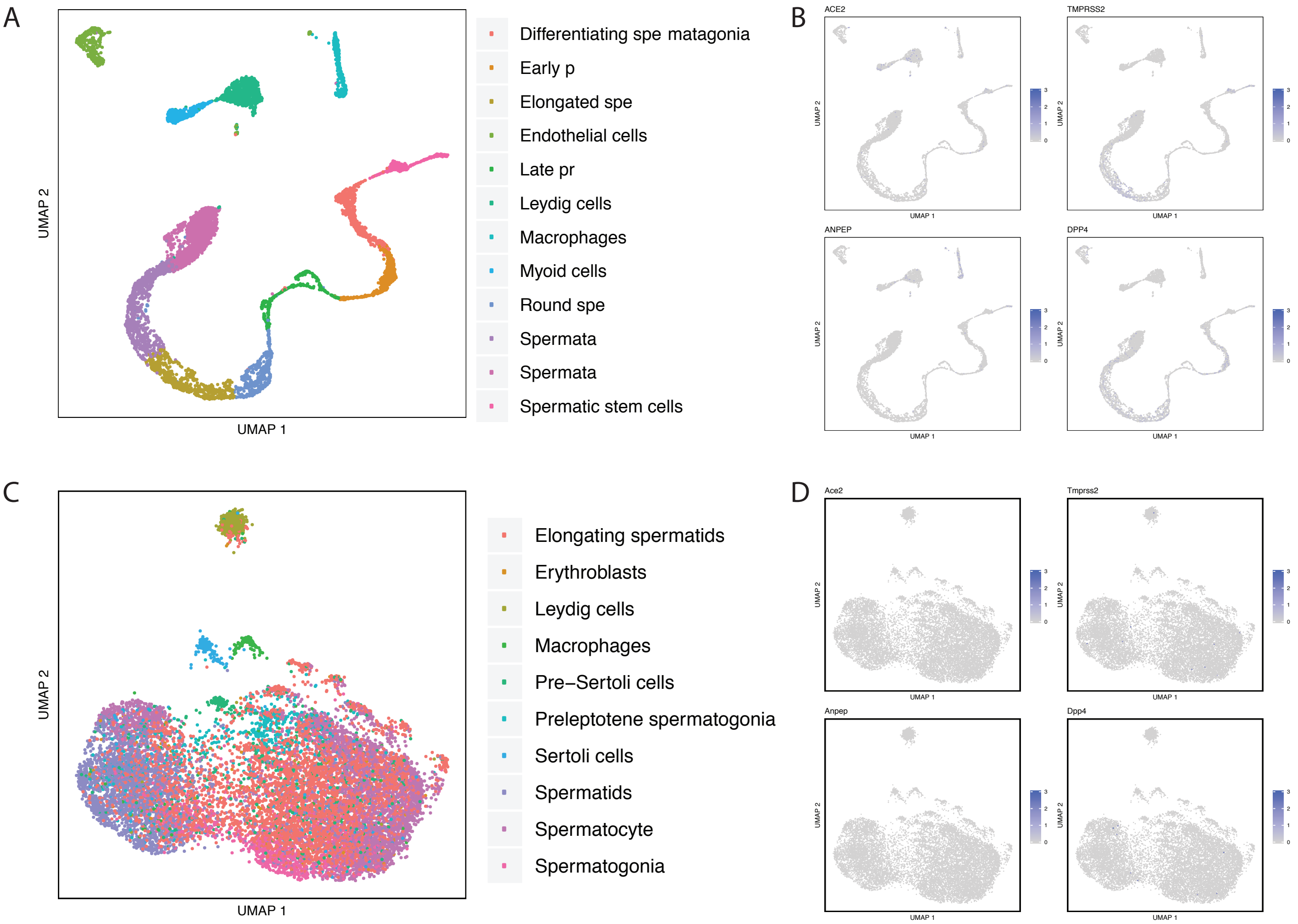


Supplemental Figure 10

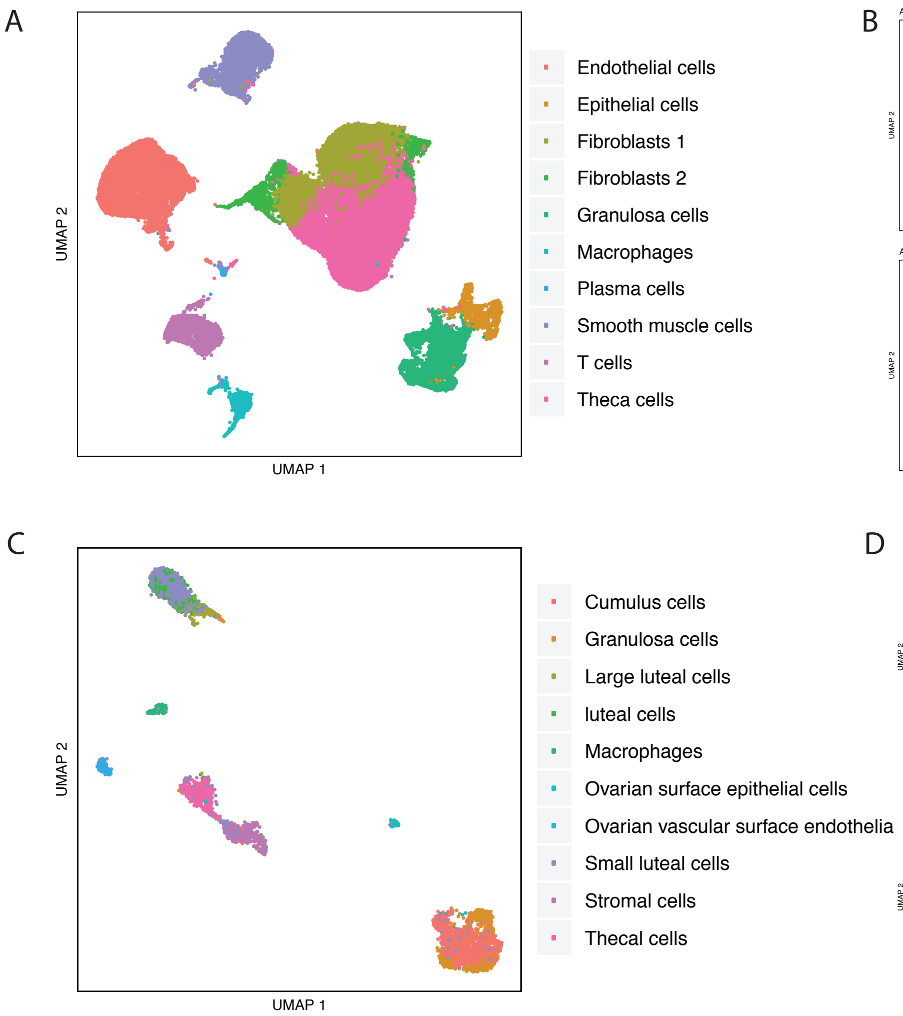


Supplemental Figure 11

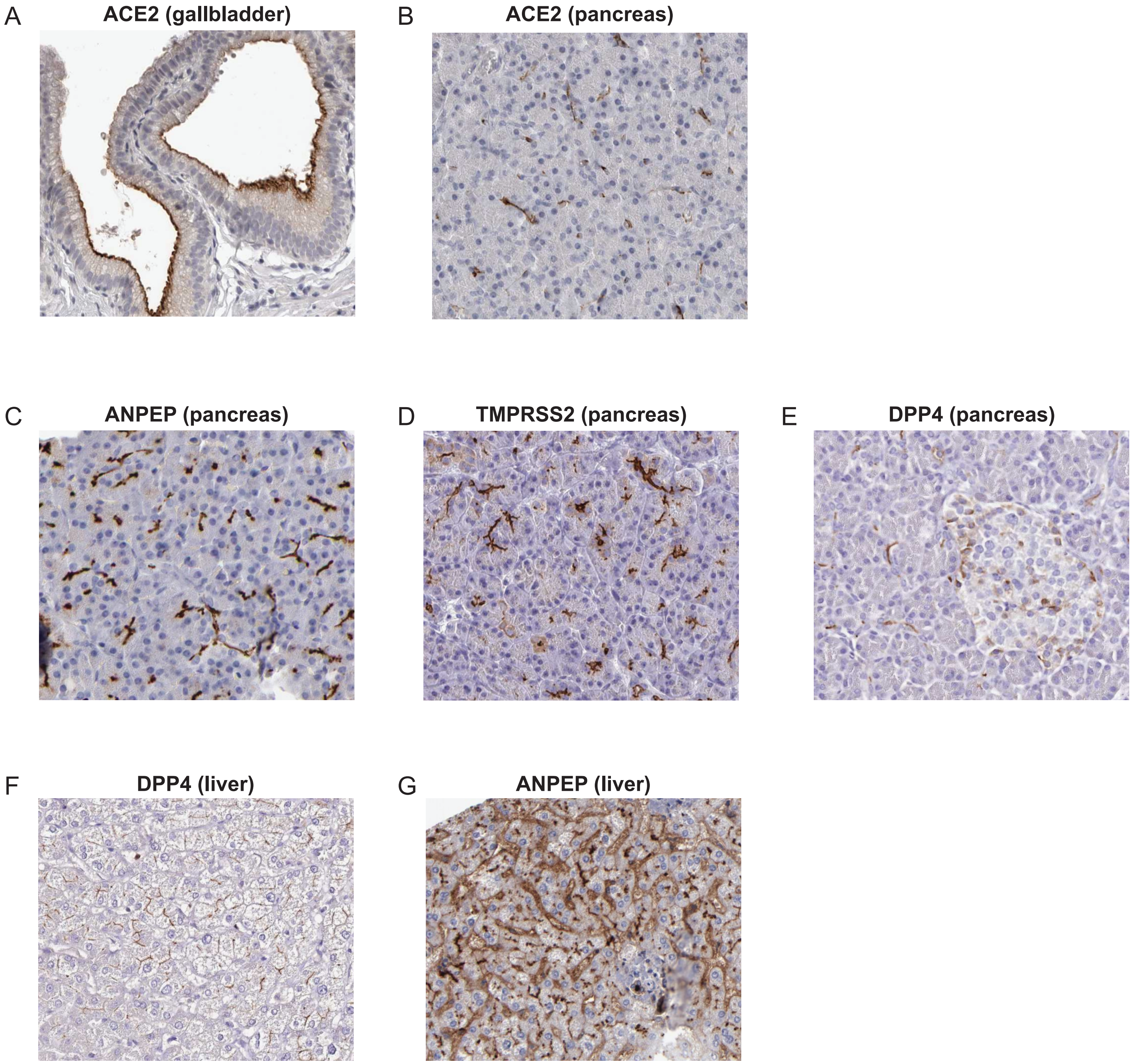


Supplemental Figure 12

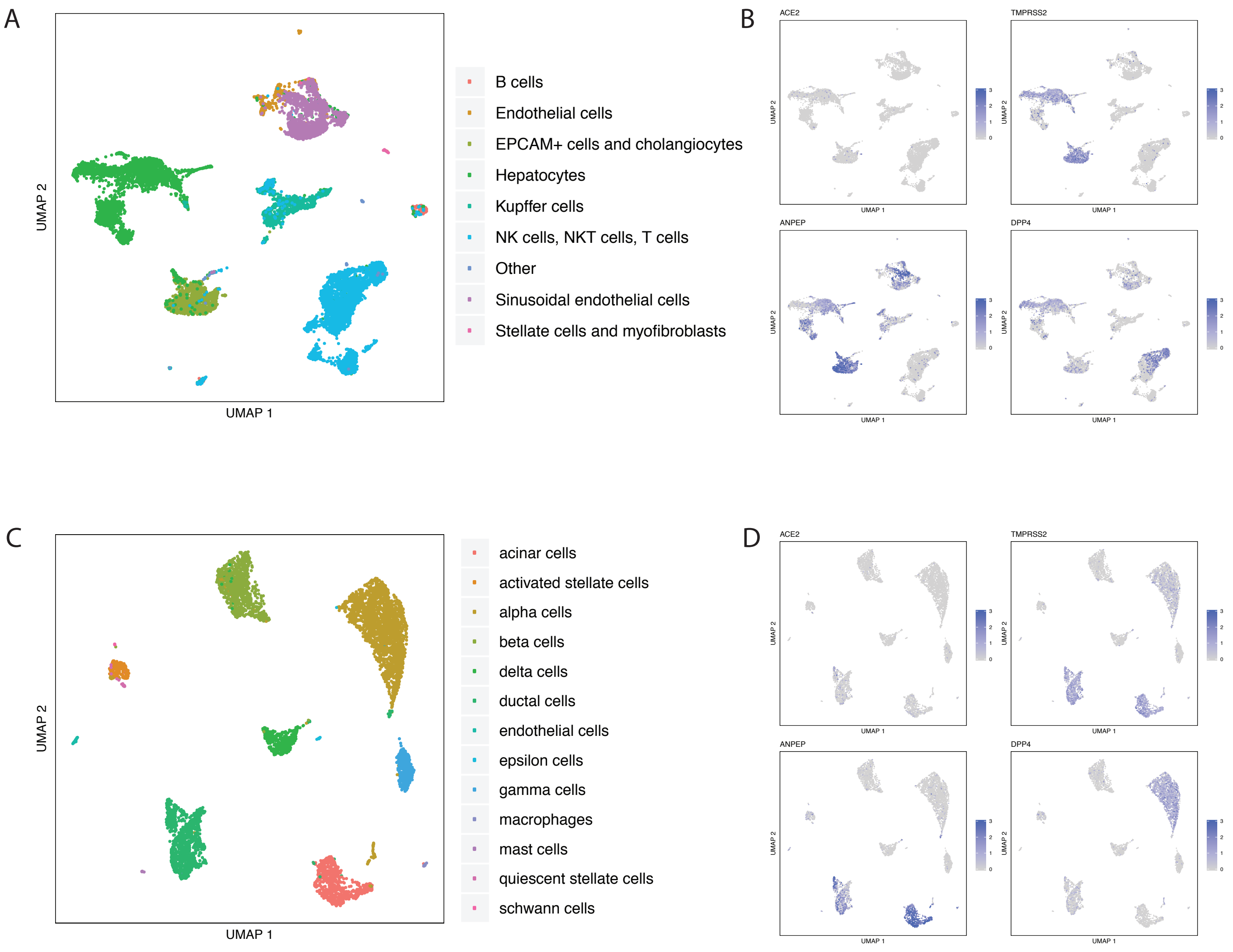


Supplemental Figure 13

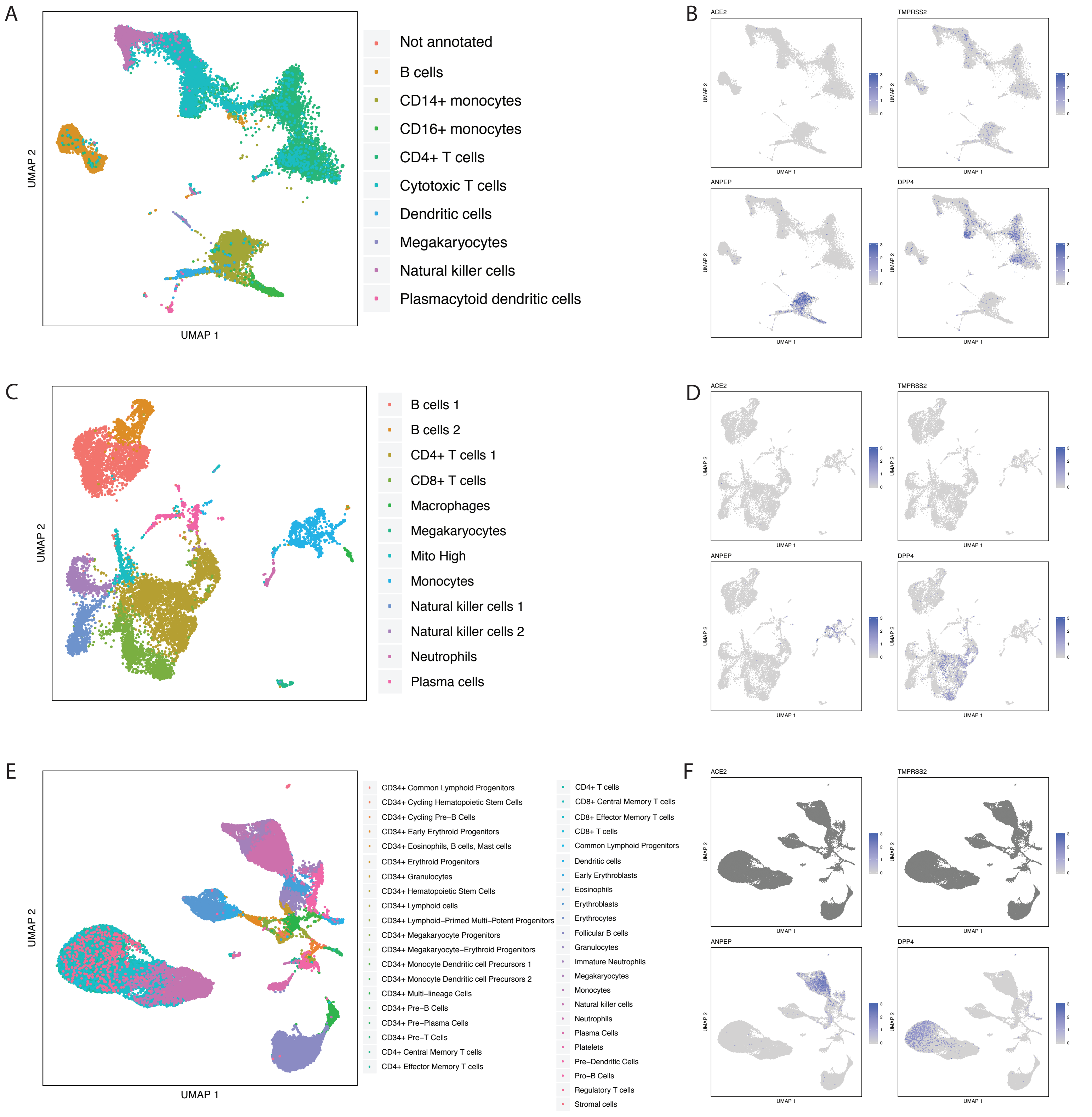


Supplemental Figure 14

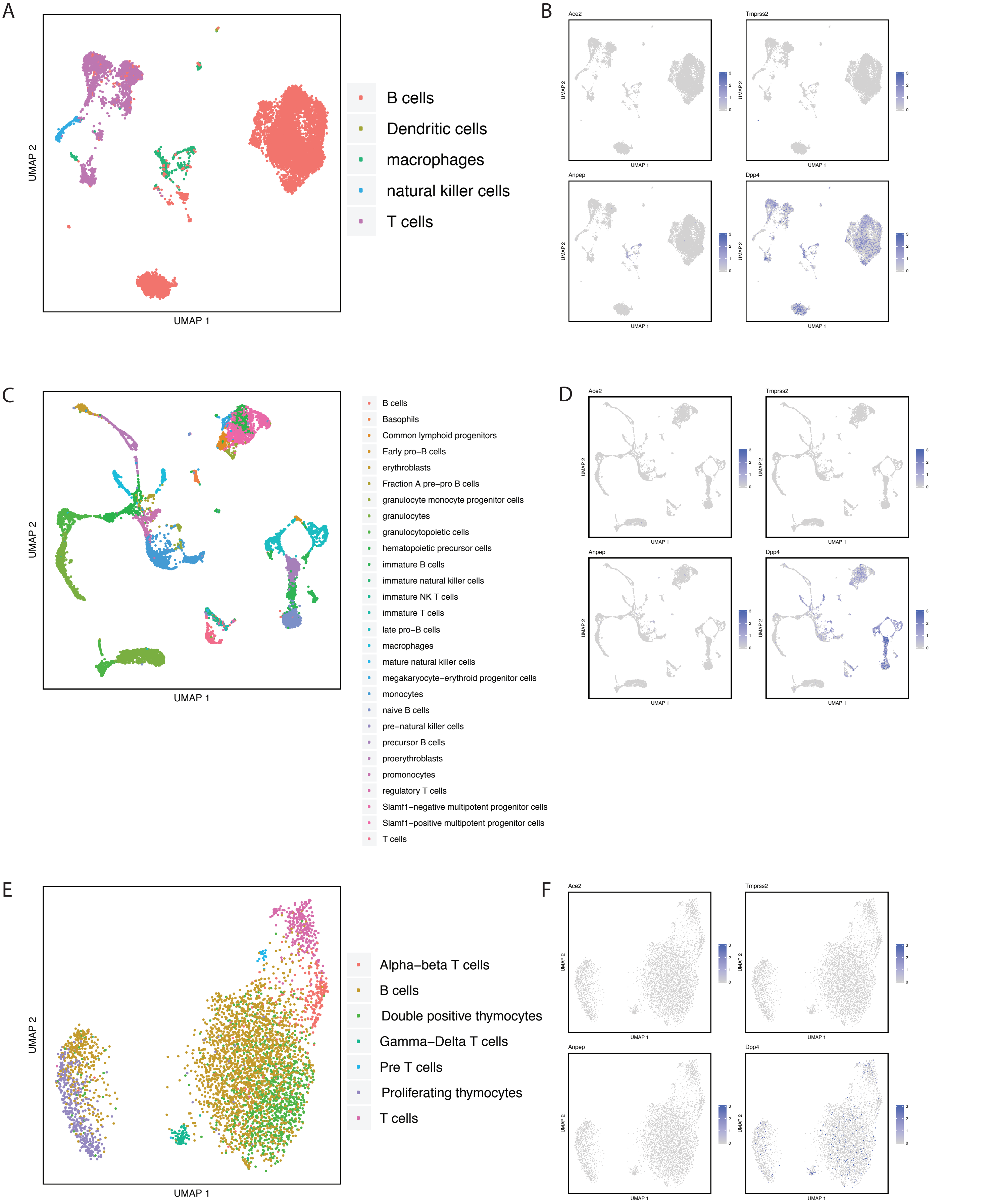


Supplemental Figure 15

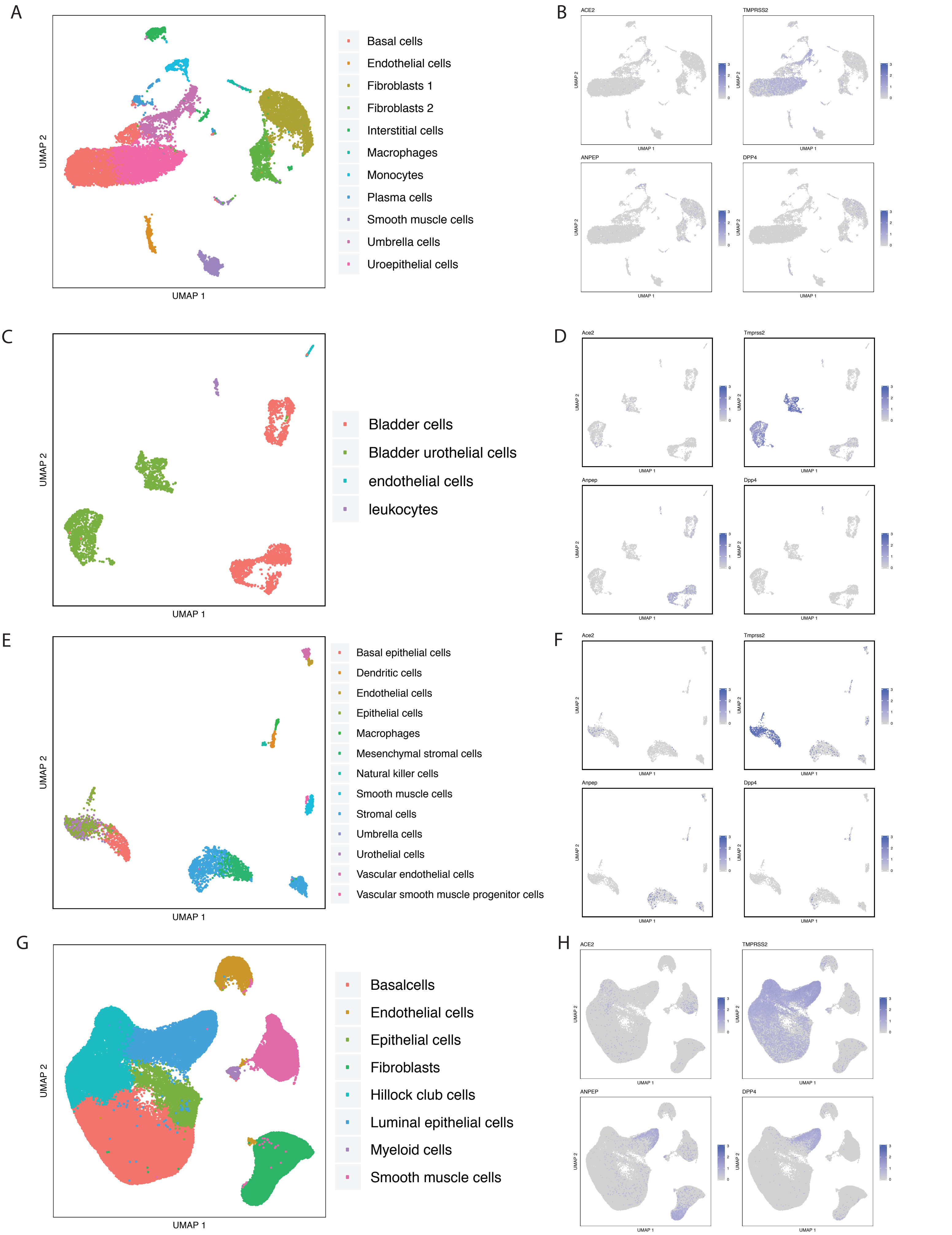


Supplemental Figure 16

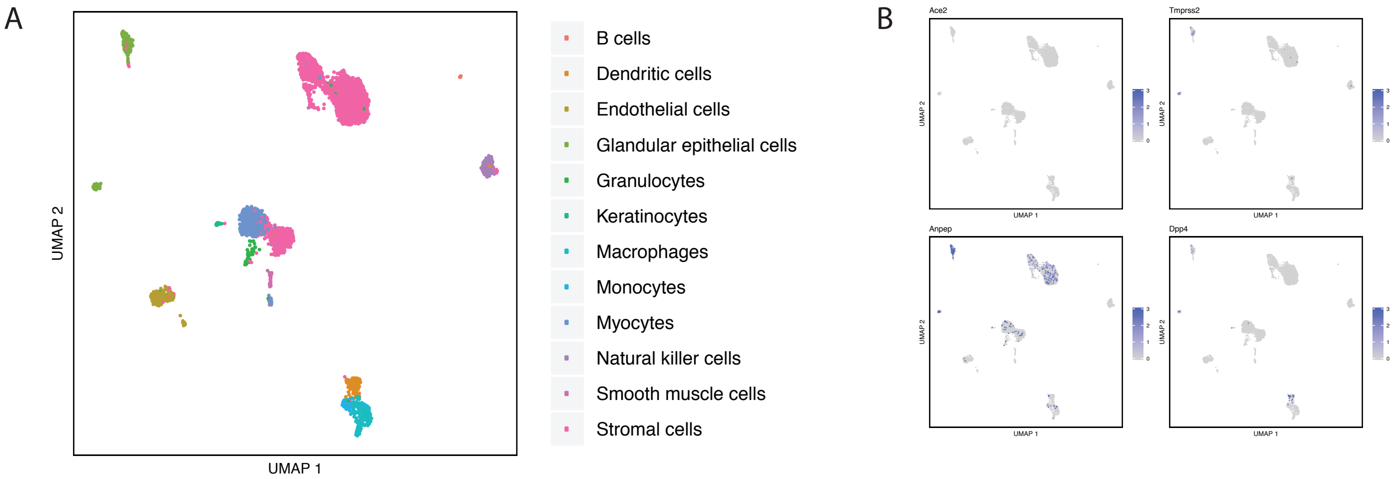


Supplemental Figure 17

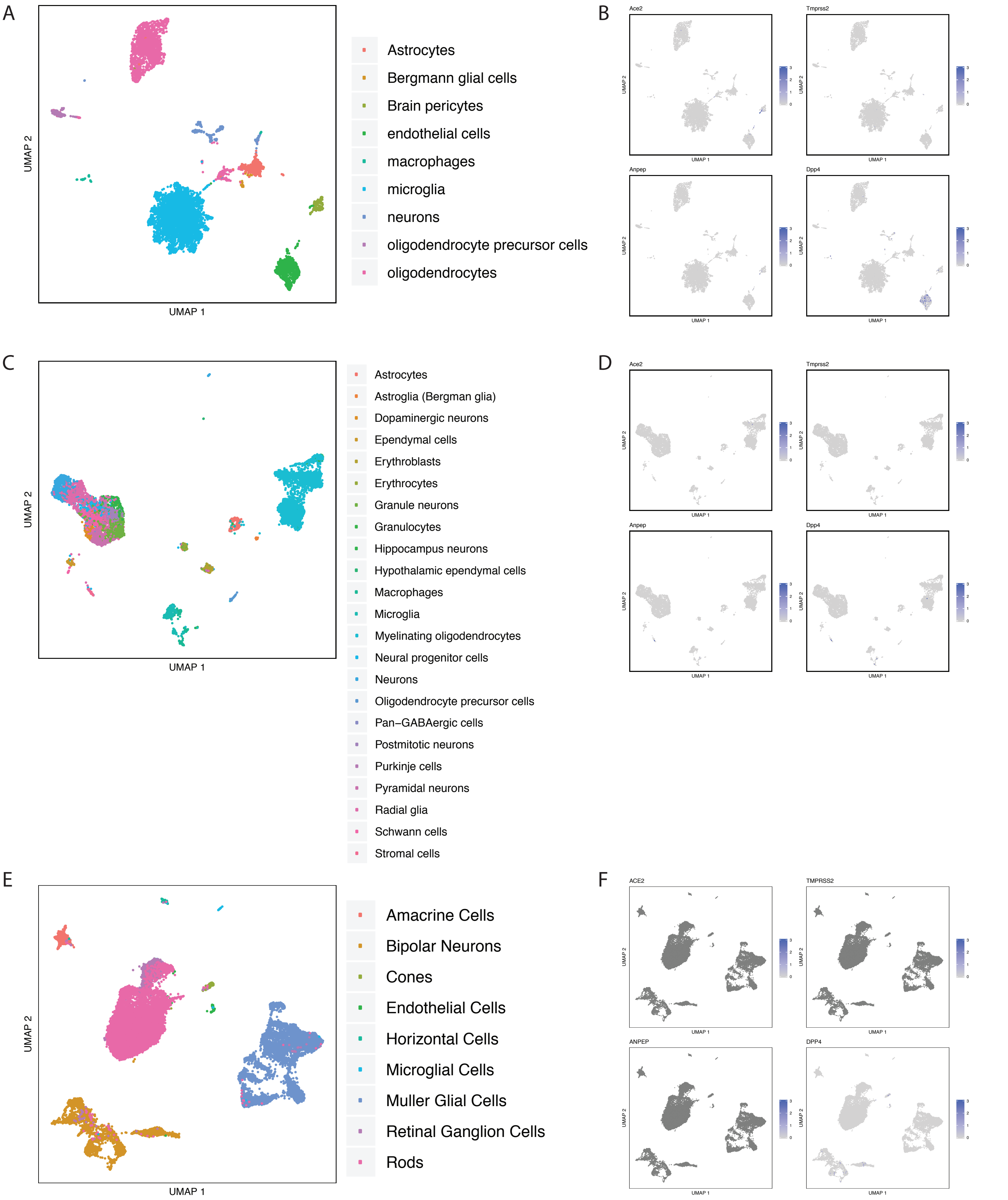


Supplemental Figure 18

#### DPP4 (brain cortex)

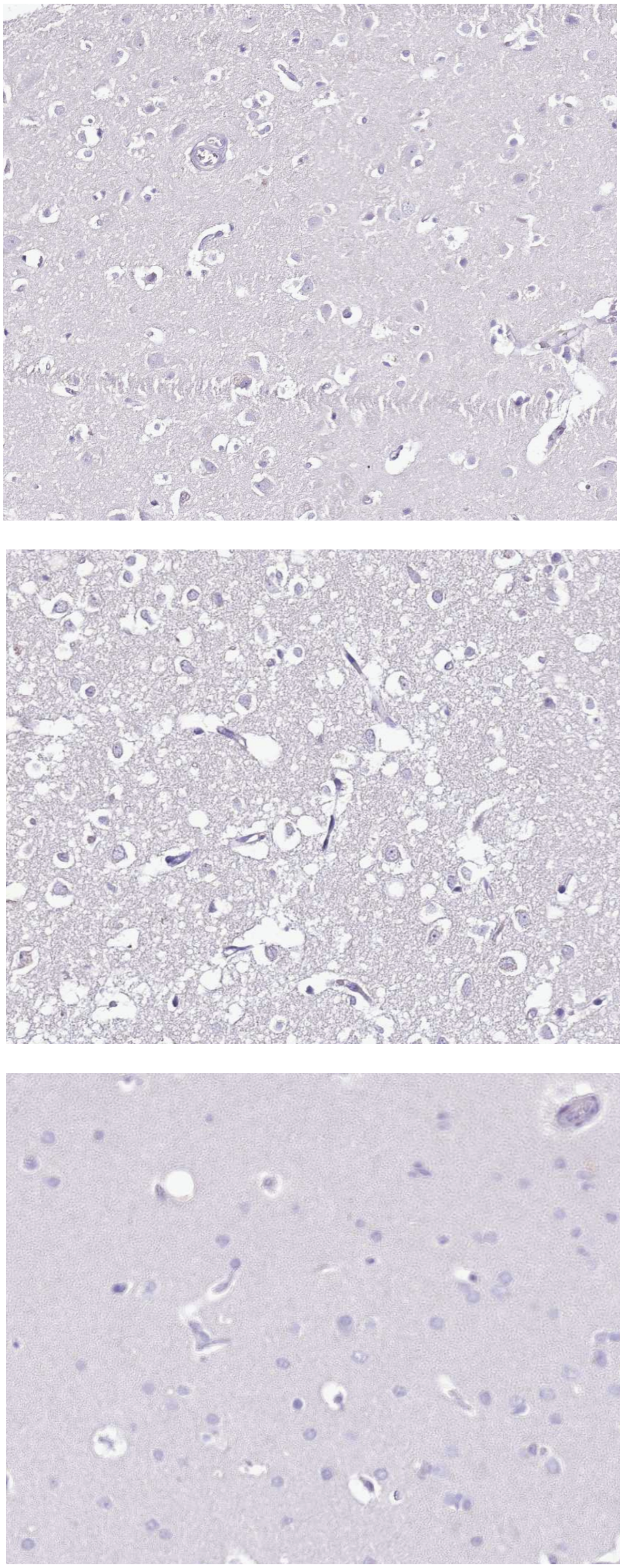

### Transcriptional Correlations to ACE2 in GTEx Lung (n = 573)

Correlation Distribution

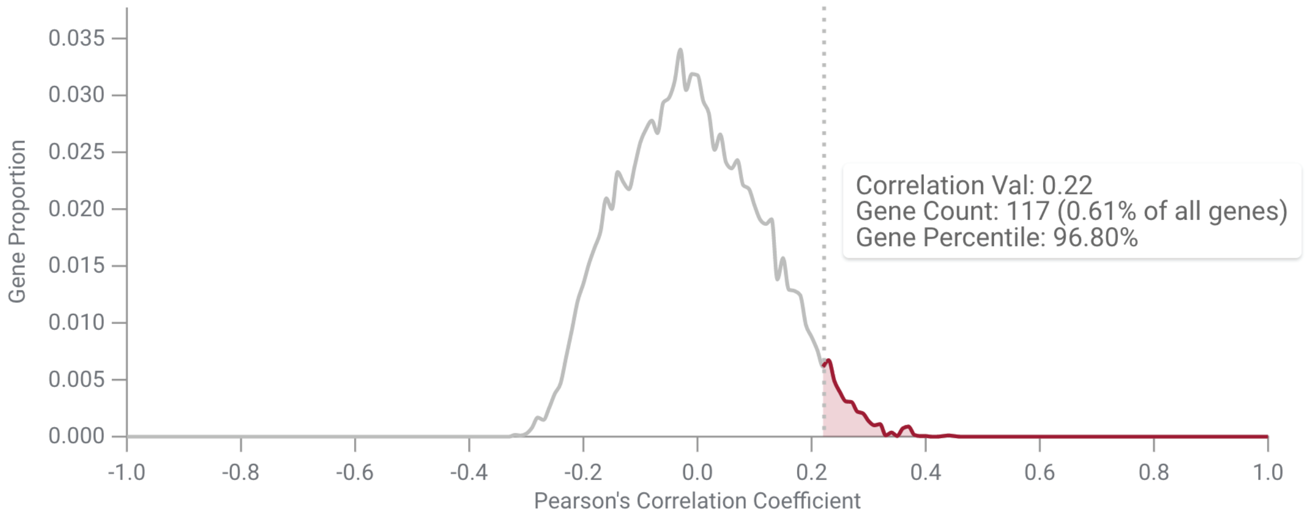


| Min | Max | | Min | Max |
|---|---|---|---|---|
| -1.00 | -1.00 | | 0.22 | 1.00 |

**Lower Range**

**Gene Count:** 0 (0.00%)

**Upper Range**

**Gene Count:** 726 (3.81%)

Correlated Genes

SFT

1-5 of 5

| | Gene | Expression Mean (TPM) | Expression Standard Deviation (TPM) | Pearson's Correlation Coefficient |
|---|---|---|---|---|
| | Spbs (SFTPB) | 5,190.10 | 21,757,558.00 | 0.29 |
| | Sp D (SFTPD) | 397.70 | 148,659.61 | 0.28 |
| | Sp G (SFTA2) | 244.91 | 26,193.68 | 0.26 |
| | Sp C (SFTPC) | 4,049.83 | 25,062,930.00 | 0.25 |
| | Sp A (SFTPA1) | 3,882.18 | 18,751,002.00 | 0.24 |
| | Nanci (SFTA3) | 114.29 | 6,650.68 | 0.23 |
| | Spa2 (SFTPA2) | 4,046.41 | 21,514,170.00 | 0.22 |

Supplemental Figure 20

#### IHC Staining of ACE2 in Nasopharynx

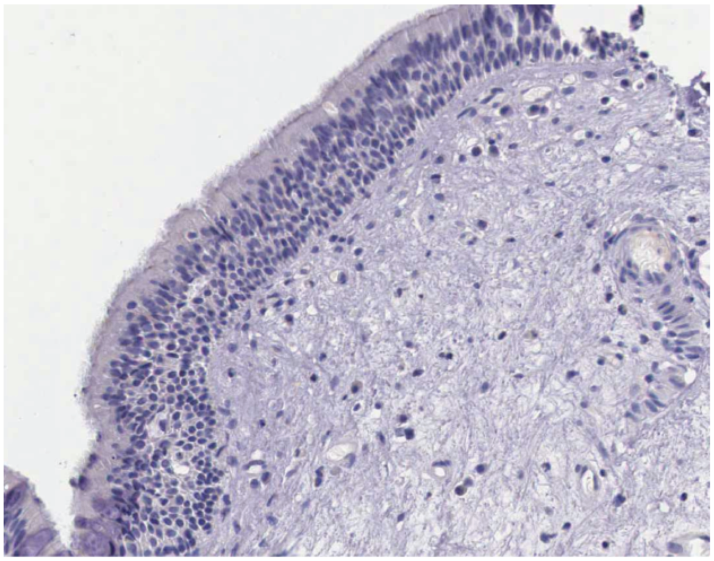

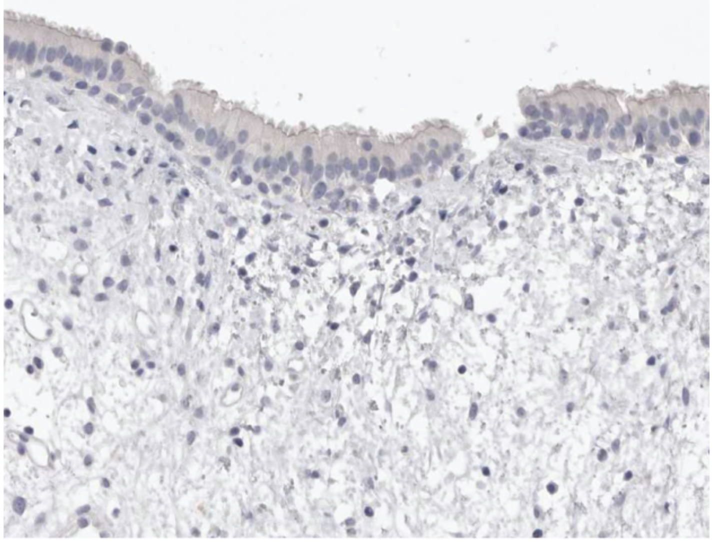

Supplemental Figure 21

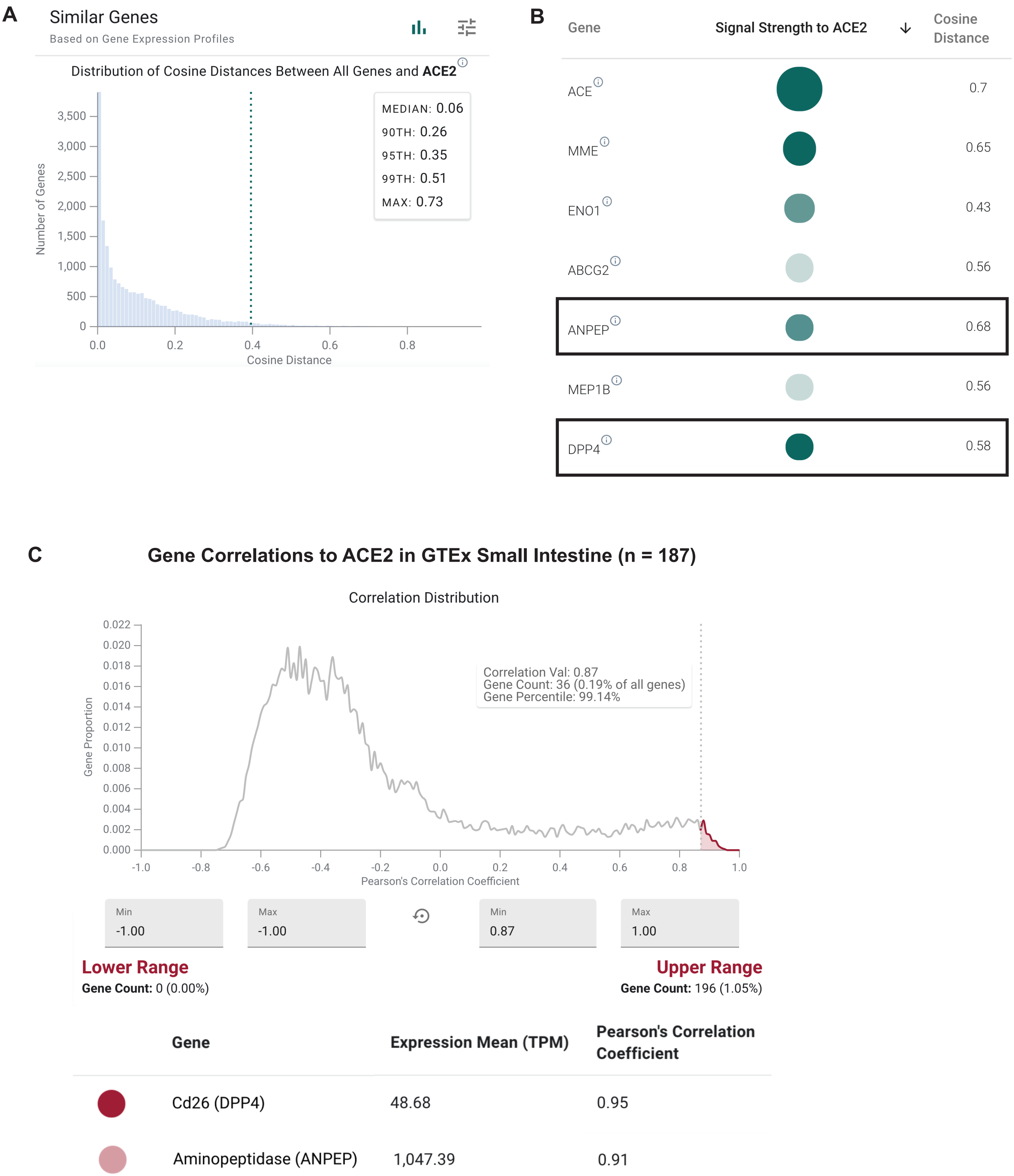


Supplemental Figure 22

A

#### Expression of Sfn in Individual Cell Populations

1 Tissue, 3 Cell Populations, 9K Cells

| Tissue | Signals Strength to Tissue | Cell | Signals Strength to Cell | Expression: ln(1 + CP10K) | Number of Cells | Percent of cells with expression > 0 | Mean ln(1+CP10K) | Cohen's D ↓ |
|---|---|---|---|---|---|---|---|---|
| tongue | | keratinocytes | | 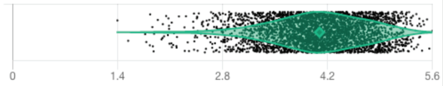 | 3406 | 100 | 4.05 | 1.15 |
| tongue | | Basal cells of epidermis | | 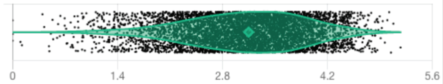 | 5510 | 99.95 | 3.08 | -0.61 |
| tongue | | Langerhans cell | | 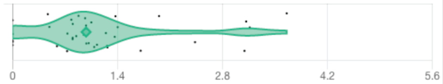 | 38 | 86.84 | 1.16 | -1.75 |

B

#### Expression of Krt6a in Individual Cell Populations

1 Tissue, 3 Cell Populations, 9K Cells

| Tissue | Signals Strength to Tissue | Cell | Signals Strength to Cell | Expression: ln(1 + CP10K) | Number of Cells | Percent of cells with expression > 0 | Mean ln(1+CP10K) | Cohen's D ↓ |
|---|---|---|---|---|---|---|---|---|
| tongue | | keratinocytes | | 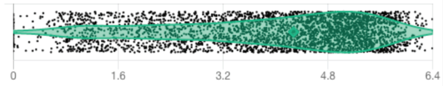 | 3406 | 99.35 | 3.9 | 1.01 |
| tongue | | Basal cells of epidermis | | 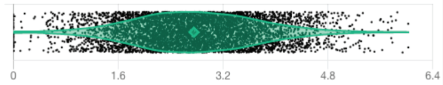 | 5510 | 99.44 | 2.78 | -0.67 |
| tongue | | Langerhans cell | | 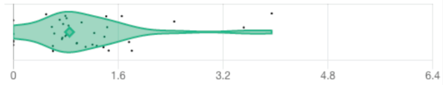 | 38 | 86.84 | 1.08 | -1.01 |

C

#### Expression of Krt10 in Individual Cell Populations

1 Tissue, 3 Cell Populations, 9K Cells

| Tissue | Signals Strength to Tissue | Cell | Signals Strength to Cell | Expression: ln(1 + CP10K) | Number of Cells | Percent of cells with expression > 0 | Mean ln(1+CP10K) | Cohen's D ↓ |
|---|---|---|---|---|---|---|---|---|
| tongue | | keratinocytes | | 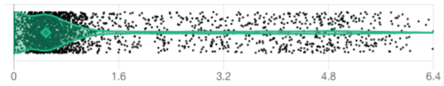 | 3406 | 75.25 | 1.04 | 0.43 |
| tongue | | Langerhans cell | | 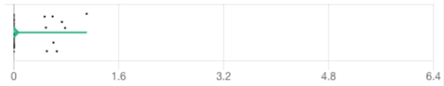 | 38 | 23.68 | 0.16 | -0.27 |
| tongue | | Basal cells of epidermis | | 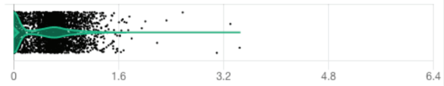 | 5510 | 49.27 | 0.3 | -0.4 |

Supplemental Figure 23

A

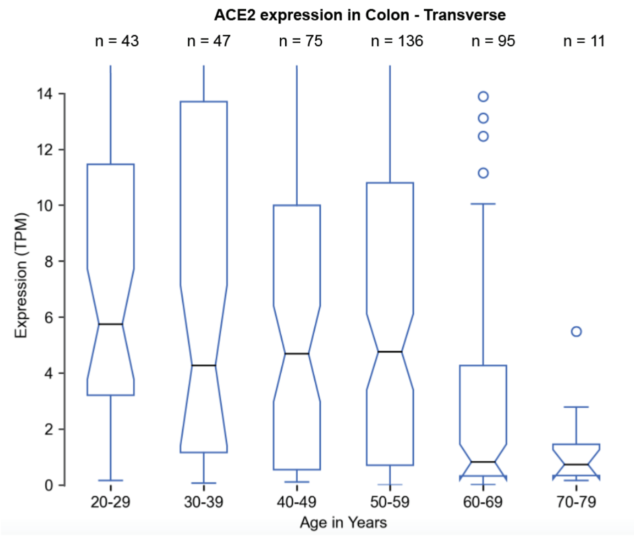


B

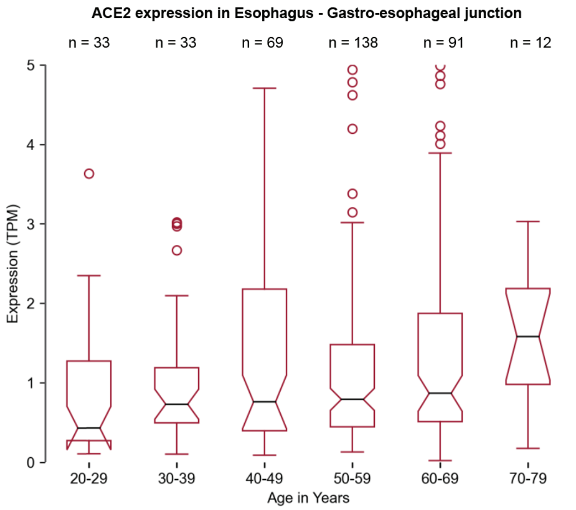


C

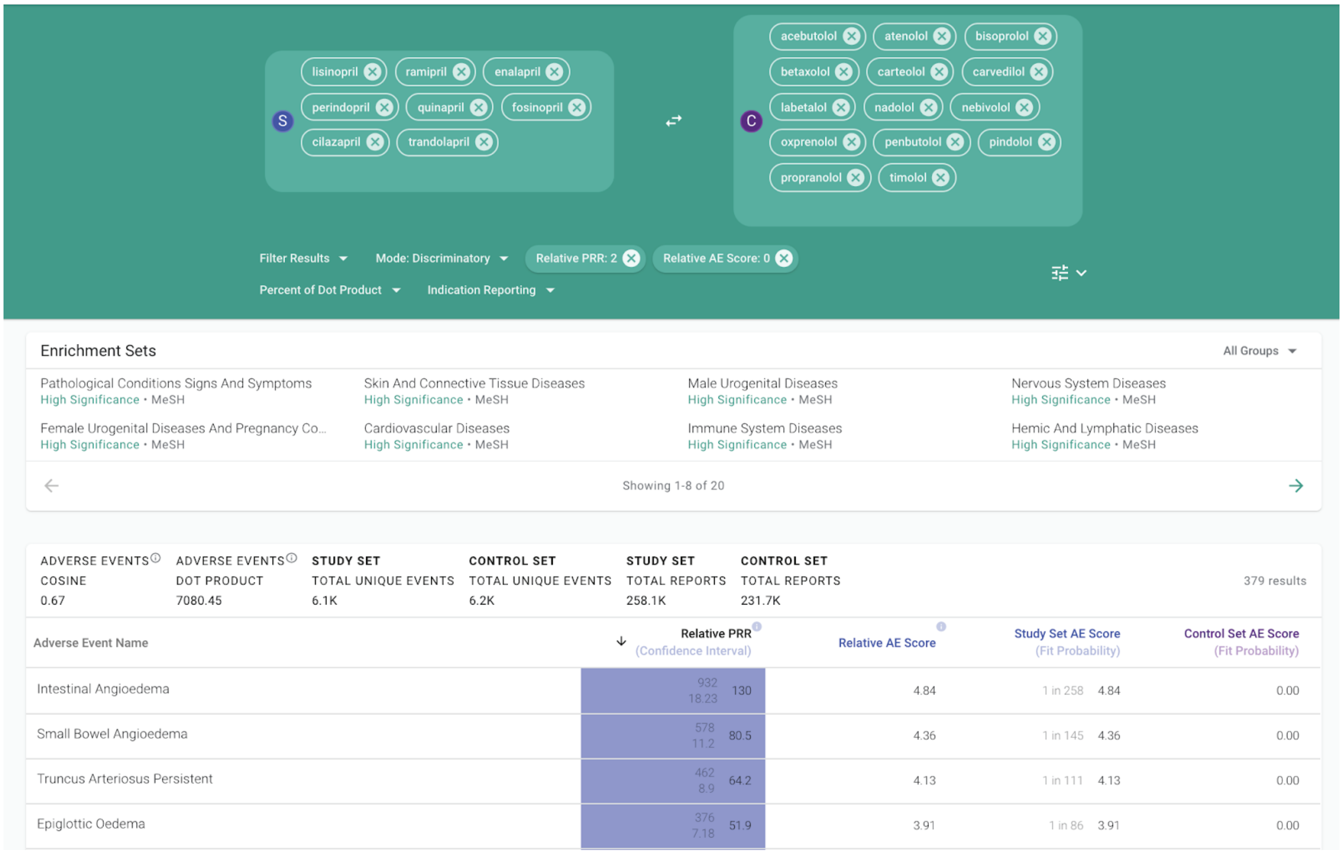


Supplemental Figure 24

#### Expression of coronavirus entry receptors in respiratory tract associated samples from GEO in comparison to GTEx Lung

##### Lung (GTEx) vs. GSE113209

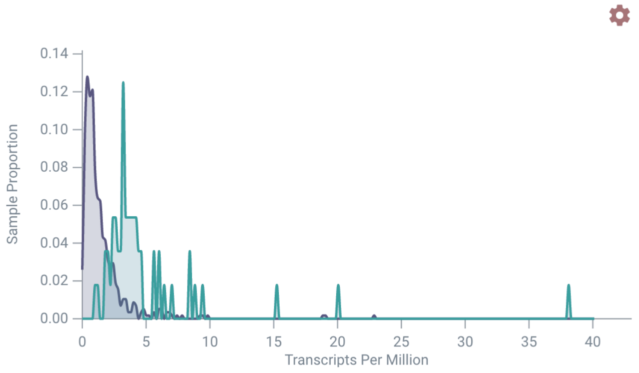


**Group A**
**Sample Count:** 578
**Mean:** 1.57 (TPM)
**Standard Deviation:** 1.96 (TPM)

**Distribution Information**
**Cohen's d:** -0.87 (Reference Group B)

**Group B**
**Sample Count:** 56
**Mean:** 5.12 (TPM)
**Standard Deviation:** 5.46 (TPM)

##### Lung (GTEx) vs. GSE124949

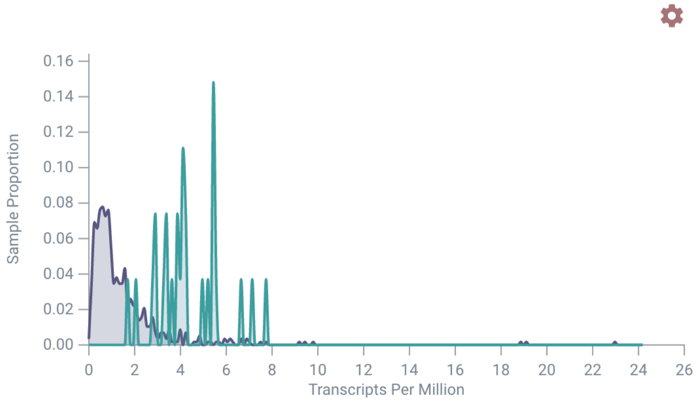


**Group A**
**Sample Count:** 578
**Mean:** 1.57 (TPM)
**Standard Deviation:** 1.96 (TPM)

**Distribution Information**
**Cohen's d:** -1.66 (Reference Group B)

**Group B**
**Sample Count:** 27
**Mean:** 4.41 (TPM)
**Standard Deviation:** 1.43 (TPM)

##### Lung (GTEx) vs. GSE97668

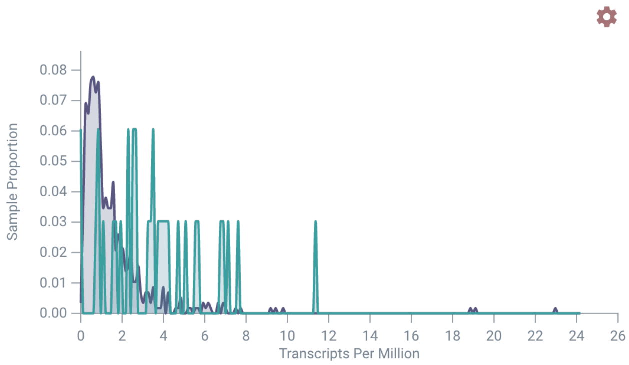


**Group A**
**Sample Count:** 578
**Mean:** 1.57 (TPM)
**Standard Deviation:** 1.96 (TPM)

**Distribution Information**
**Cohen's d:** -0.93 (Reference Group B)

**Group B**
**Sample Count:** 33
**Mean:** 3.63 (TPM)
**Standard Deviation:** 2.45 (TPM)

##### Lung (GTEx) vs. GSE93526

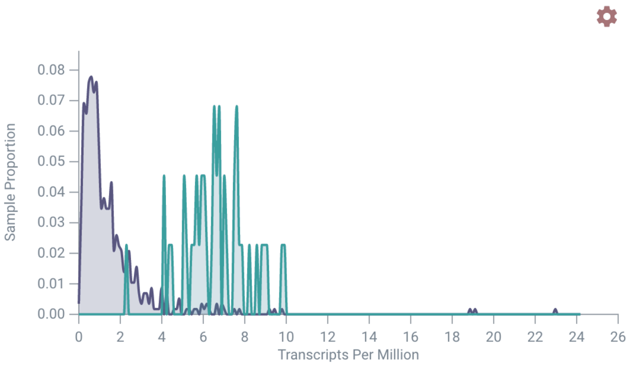


**Group A**
**Sample Count:** 578
**Mean:** 1.57 (TPM)
**Standard Deviation:** 1.96 (TPM)

**Distribution Information**
**Cohen's d:** -2.88 (Reference Group B)

**Group B**
**Sample Count:** 44
**Mean:** 6.67 (TPM)
**Standard Deviation:** 1.56 (TPM)

Supplementary Figure 25
